# Longitudinal DNA methylation dynamics distinguish Persistent and Remitted ADHD in childhood and adolescence

**DOI:** 10.1101/2025.10.21.681830

**Authors:** Jo Wrigglesworth, Valentine Chirokoff, Peter D. Fransquet, Jeffrey M. Craig, Tim J. Silk

## Abstract

Attention-deficit/hyperactivity disorder (ADHD) is a neurodevelopmental disorder, with symptoms that remit or persist over time. Biological mechanisms underlying symptom change remain poorly understood, though epigenetic processes, like DNA methylation (DNAm), may serve as dynamic biomarkers of clinical outcomes. We examined change in DNAm in children and adolescents with and without ADHD, including differences between remitted and persistent ADHD.

**Methods:** We analyzed 219 saliva samples from 94 participants (aged 9.5-14.5 years) attending 2 to 3 waves of NICAP. DNAm was profiled using the Infinium MethylationEPIC BeadChip. Linear mixed models adjusted for age, sex, medication, batch and cell-proportion assessed longitudinal change. Comparisons included (1) all ADHD cases versus controls, (2) persistent or remitted ADHD versus controls, and (3) persistent versus remitted ADHD. False discovery rate correction controlled for multiple testing (FDR p<0.05).

**Results:** No CpGs were statistically different between ADHD and controls, after correction. Compared to controls, the average DNAm at 5 sites was statistically different in persistent or remitted ADHD, and methylation of 3 additional CpGs differentially changed over time in persistent ADHD. Remitted differed from persistent ADHD at four CpGs, including cg21443143 (*ZFAT*), which showed a unique decline over time. Gene enrichment links findings to brain structures and function, though these did not survive multiple testing.

**Conclusion:** Our study identified several epigenetic differences between remitted and persistent ADHD outcomes from typical development, and from each other. Given the early stage of this research, our findings warrant further prospective epigenome-wide studies into these diagnostic trajectories.

## INTRODUCTION

Attention-deficit/hyperactivity disorder (ADHD) is a common neurodevelopmental disorder, affecting ∼5% of children and adolescents worldwide (1). Symptoms of inattentiveness and/or hyperactivity and impulsivity vary within and between individuals (2), and can persist into adulthood (3). The factors underlying variation in symptom outcomes remain incompletely understood, though both genetic and environmental contributors have been reported (4–6)

Epigenetic processes, which regulate gene expression in response to genetic and environmental influences, have attracted increasing attention in ADHD research (7). The most widely studied epigenetic mechanism is DNA methylation (the addition of a methyl molecule to a cytosine of a cytosine-phosphate-guanine ‘CpG’ dinucleotide), which has been linked to brain development and neurodevelopmental disorders (8, 9).

A growing number of epigenome wide association studies (EWAS) have investigated DNA methylation in ADHD (10–20). Most of these are cross-sectional studies investigating methylation differences between ADHD and controls, and are often limited to childhood samples. Moreover, findings are inconsistent across probes, and are not always corrected for multiple comparisons

Of the few longitudinal studies of ADHD to date, most have related early life DNA methylation to later childhood or adolescent symptoms (13, 14, 18), and therefore do not directly inform on patterns of epigenetic change. Only one case-control study has compared DNA methylation between individuals with persistent or remitted ADHD, but this focused on young adults (mean age of 20.9 years) (21).

Our primary aim provides contextual information about general ADHD-related dynamics in methylation by examining whether longitudinal changes in DNA methylation differ between children and adolescents with and without ADHD.

Our secondary, and clinically relevant aim determines whether DNA methylation change distinguishes remitted from persistent ADHD. This work provides critical evidence towards the development of future prognostic tests that can operate within the ADHD population, rather than through case-control contrasts. Because epigenetic mechanisms are dynamic and potentially modifiable, identifying such signatures may improve prediction of clinical outcomes, provide tools for monitoring and guiding treatment, and biological pathways underlying ADHD symptom change.

## METHODS

### Study participants

Data for this study comes from the Neuroimaging of the Children’s Attention Project (NICAP) (22), a sub-study of the Children’s Attention Project (CAP) (23). Recruitment into CAP has been described in detail elsewhere (23). Briefly, typically developing children (aged 6-8 years) were recruited from 43 socio-economically diverse primary schools across the Eastern and Western Melbourne metropolitan regions, Australia. CAP initially screened for ADHD symptoms using independent parent and teacher reports on the Conners 3 ADHD index (24), and confirmed ADHD diagnoses using the NIMH Diagnostic Interview Schedule for Children-IV (DISC-IV) (25).

NICAP recruited CAP participants at the 36-month follow-up assessment of the study, when children were 10 years of age (26). Two additional visits were conducted at 18-month intervals, when participants were aged 11.5 and 13 years (26). This repeated-measures design provided DNA methylation data at multiple time points for each participant, therefore enabling longitudinal analysis of epigenetic change within individuals. Diagnostic assessment for ADHD was repeated at the first and last NICAP waves.

Children were classified as the ADHD group if they had met criteria for ADHD at the first wave of epigenetic sampling, or three years earlier when participants were recruited into CAP. To reflect the changing nature of ADHD symptoms, cases were further categorized by diagnostic outcome at the final NICAP wave: participants meeting criteria were defined as persistent ADHD, and those no longer meeting criteria were defined as remitted ADHD. Controls did not meet criteria for ADHD at any wave. Control comparisons were included to provide context on general ADHD-related methylation dynamics, while clinically relevant contrasts were within ADHD groups (persistent vs. remitted) for potential prognostic insight.

Children were eligible for this study if they had both a saliva sample and diagnostic assessment from at least two waves. In total, 219 saliva samples were collected from 94 participants: 63 attended two waves, and 31 attended three waves of assessment. The distribution of chronological age at each wave of assessment are presented in **Figure 1**.

**Figure 1.**
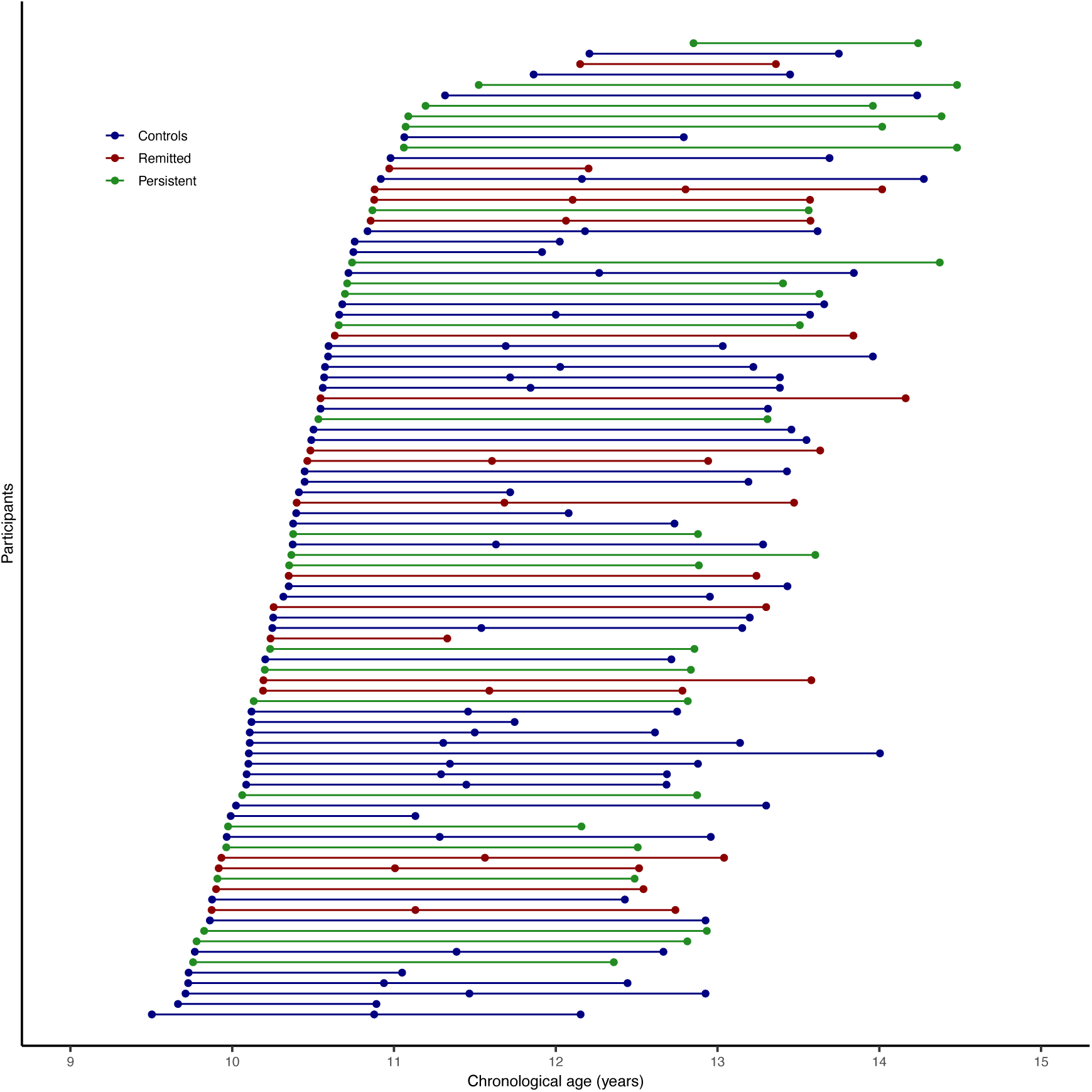
Distribution of participant ages at each NICAP wave where saliva samples were collected for DNA methylation analysis. Points are colored by follow-up diagnostic status: blue = controls, red = remitted ADHD, green = persistent ADHD. Horizontal spread is jittered for visibility.

This study was approved by the Royal Children’s Hospital Human Research Ethics Committee, Melbourne, Australia (#34071). Written informed consent was obtained from the parents.

### DNA methylation pre-processing and quality control

Genomic DNA was isolated from ∼3 mL saliva samples that were collected at each assessment via passive drool into a 50 mL centrifuge tube (Eppendorf South Pacific, NSW, Australia) and stored at -80° C until analysis. DNA samples were bisulphite treated and hybridized to Infinium MethylationEPIC BeadChip arrays, version 1.0 (Illumina, San Diego, CA, USA), at NTX-Dx (Diagenode, Ghent, Belgium). These arrays generate data from over 850,000 CpGs across the human genome and are enriched for probes in regulatory regions.

Quality control was performed in R using the R *minfi* package (27). Probes were mapped to genomic features of the human genome assembly GRCh37 (hg19) using *IlluminaHumanMethylationEPICanno.ilm10b4.hg19*. Data across all visits were normalized collectively using subset quantile normalization (28, 29). Probes were removed if they had a poor-quality signal (113,011 with detection p>0.01); were on a sex chromosomes (n=16,376); overlapped a single nucleotide polymorphisms (SNP) at the CpG or extension site (n=26,455), or mapped to multiple locations (n=34,471) (30). Beta-values represent the proportion of methylation at a CpG site (methylated signal intensity divided by the total intensity), and M-values, the log^2^ ratio of methylated to unmethylated signal intensities. Both measures were calculated for the 675,546 remaining probes, but M-values were used for statistical analysis (31).

Cell-type heterogeneity was addressed using EpiDISH (32). Because buccal samples contain negligible fibroblasts, only epithelial and leucocyte proportions were relevant, and these are inversely proportional. We therefore included epithelial proportion as a covariate, which is mathematically equivalent to adjusting for leucocyte proportion.

### CpG selection

Given the longitudinal nature of the data, we focused on probes showing DNA methylation change within individuals over time, rather than static cross-sectional differences. To address Aim 1 (longitudinal differences between ADHD and controls), the 675,546 quality-controlled probes were filtered for those with significant change over time (nominal p < 0.05) and had a change of more than 5% in the full sample (**Figure 2**). Change at each probe was modelled using unadjusted linear mixed models (LMM) with participant as a random intercept.

**Figure 2.**
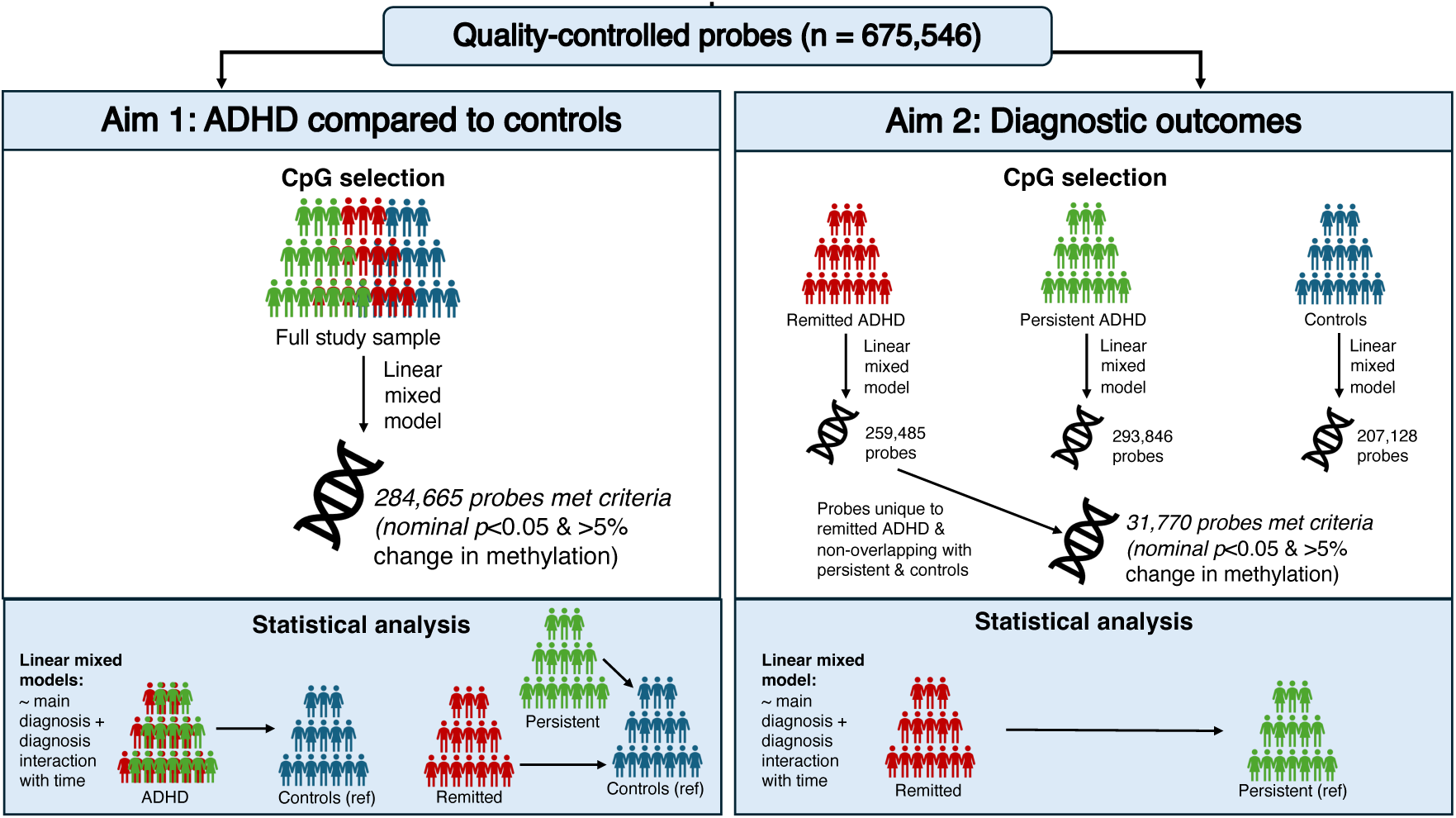
Flow diagram showing the selection of CpG sites exhibiting longitudinal DNA methylation change. Probes passing quality control and longitudinal change criteria (p <0.05 and > 5% change) were filtered to identify those unique to diagnostic groups. The diagram summarizes filtered steps, participant numbers (94 total, 63 with 2 waves, 31 with three) and the comparison performed (ADHD vs. controls; persistent vs. remitted ADHD).

To determine whether longitudinal change in DNA methylation was specific to diagnostic outcomes (persistent vs. remitted), the same criteria (p <0.05 and > 5% change) were applied to group-specific LMM. Probes unique to remitted ADHD (i.e., meeting criteria in remitted ADHD, and not overlapping with controls or persistent ADHD) were selected for further analysis (**Figure 2**).

When individual probes are cited in the Results, both the probe name and the relevant annotated gene(s), if present, are provided (e.g., cg12881363 – *MOV10L1*, cg02208776 – *HSPE1; HSPD1*, and cg11124426).

### Gene enrichment analysis

Gene enrichment analyses were performed on the larger sets of probes meeting our criteria for longitudinal change, not limited to only the final diagnostic-group-significant CpGs, to explore their biological context. Genes annotating to selected probes were tested for tissue enrichment using RNA-seq data from the Human Protein Atlas (HPA) (33) via the *TissueEnrich* package (34), then for enrichment in 47 brain tissue regions (16 brain regions, plus their super-structures, which are structures inherited through a hierarchical brain organization e.g., M1 primary motor cortex inherits super-structures like the cerebral cortex and frontal neocortex) across the developmental stages (prenatal; 0-2 years; 3-11 years; 12-19 years, and >19 years) by the Allan Brain Atlas (ABA), using the *ABAenrichment* package (35).

Functional enrichment of differentially methylated probes in molecular, cellular and biological processes and pathways was tested using Gene Ontology (GO) (36, 37) and the Kyoto Encyclopedia of Genes and Genomes (KEGG) databases, respectively (38). Here we used the *gometh* function from the *missMethyl* package (39) as this adjusts for probe number and multiple annotations.

Differentially methylated probes were further examined for gene-disease associations recorded using ClinVar (40), ClinGen (41) and PsyGeNET (42) databases via *disgenet2r* (43). Multiple comparisons were controlled at a false discovery rate (FDR) of 5% (44), except for ABA enrichment, which used the package’s default minimum family-wise-error rate (FWER).

### Statistical analysis

Differential DNA methylation was analyzed using R v4.4 (45) *limma* package. Longitudinal differences in DNA methylation were first assessed between ADHD and controls using adjusted LMM, which regress selected probes (meeting longitudinal change criteria, refer to section ‘CpG Selection’) on the main effect of diagnosis and its interaction with time. Comparisons included: (1) all ADHD cases vs. controls, (2) persistent or remitted ADHD vs. controls, and (3) persistent vs. remitted ADHD. Models were adjusted for sex, medication use, batch, and epithelial cell proportion, with participant as a random intercept. Because the clinical application of this study is prognostic rather than diagnostic, we prioritized persistent vs. remitted ADHD comparisons, with ADHD-control comparisons included for context. We correct for multiple comparisons using a Benjamini-Hochberg FDR corrected p-value less than 5% (44).

For probes significantly differing between persistent and remitted ADHD, residual change scores were used to further examine associations between DNA methylation change and ADHD symptom change. Residual change in methylation was defined by regressing M-values at last follow-up on baseline values, adjusted for baseline age, batch, epithelial cell proportion and time interval. Residual change in symptoms was similarly defined using parent-reported Conners 3 ADHD Index (24), adjusted for baseline age and time interval. For both measures, a positive residual change score represents a higher level of methylation or symptom severity at follow-up, after considering data collected at baseline. Associations between methylation change and symptom change were tested with linear regression models adjusted for sex and medication use.

## RESULTS

### Study participants

We analyzed 219 saliva samples from 94 participants, 47% of whom had a childhood history of ADHD (**Table 1**). Among the 44 ADHD cases, 57% retained a diagnosis at their final wave (persistent ADHD), while 43% no longer met criteria (remitted ADHD). Across waves and ADHD groups, males represented more than half of the eligible participants.

**Table 1.** Characteristics of study cohort. Demographic and clinical characteristics of participants at each wave of the NICAP study. Values show numbers of participants (n) and percentages (%) or mean values with ranges, stratified by diagnostic status: controls, remitted ADHD and persistent ADHD. ADHD symptom severity was measured using parent-reported Conners 3 ADHD Index scores. Medication use reflects any prescribed behavioral medication at each wave

| Characteristic | Controls | Remitted ADHD | Persistent ADHD | Total |
| --- | --- | --- | --- | --- |
| n (% male) |  |  |  |  |
| Wave 1 | 50 (70%) | 18 (72%) | 24 (71%) | 92 (71%) |
| Wave 2 | 32 (100%) | 12 (83%) | 1 (100%) | 45 (96%) |
| Wave 3 | 40 (63%) | 17 (71%) | 25 (72%) | 82 (67%) |
| Chronological age mean (range) |  |  |  |  |
| Wave 1 | 10.4 (9.5-11.9) | 10.4 (9.9-11.0) | 10.5 (9.8-11.5) | 10.4 (9.5-11.9) |
| Wave 2 | 11.7 (10.9-13.4) | 11.8 (11.0-12.8) | 12.9 | 11.7 (10.9-13.4) |
| Wave 3 | 13.2 (12.2-14.3) | 13.3 (12.5-14.2) | 13.3 (12.2-14.5) | 13.3 (12.2-14.5) |
| ADHD symptoms, mean (range) |  |  |  |  |
| Wave 1 | 1.0 (0.0-7.0) | 7.8 (0.0-17.0) | 11.0 (0.0-20.0) | 4.8 (0.0-20.0) |
| Wave 2 | 1.2 (0.0-5.0) | 6.8 (1.0-17.0) | 13.0 | 3.0 (0.0-17.0) |
| Wave 3 | 0.4 (0.0-7.0) | 5.1 (0.0-15.0) | 10.5 (0.0-19.0) | 4.3 (0.0-19.0) |
| Medication use, n (%) |  |  |  |  |
| Wave 1 | 0 (0.0%) | 4 (22%) | 8 (36%) | 12 (14%) |
| Wave 2 | 0 (0.0%) | 4 (33%) | 1 (100%) | 5 (11%) |
| Wave 3 | 0 (0.0%) | 2 (13%) | 7 (32%) | 9 (13%) |

Within the ADHD group, 20 participants reported prescribed medication use for behavior at one or more visits. These included methylphenidate (n=8), atomoxetine (n=2), lisdexamfetamine (n=3), fluoxetine (n=2), risperidone (n=1), or combinations (i.e., risperidone + atomoxetine [n=1], clonidine + methylphenidate [n=1], fluoxetine + lisdexamfetamine [n=1], or dexamphetamine + lisdexamfetamine [n=1]). Only one remitted ADHD participant used medication (methylphenidate) at all visits; the others used mediation intermittently.

### CpG selection for longitudinal change and diagnostic outcomes

Out of the 675,546 probes surviving quality control, 284,665 (42%) showed significant change in methylation across development, comprising 145,493 hypomethylated and 139,172 hypermethylated sites (**Supplementary Figure 1a**). Among these, 188,942 probes mapped to a single gene which were predominantly expressed in brain tissue (**Supplementary Figure 1b**), and were enriched in seven brain regions in childhood (0-2 years and 3-11 years of the ABA data, all FWER adjusted p<0.05; **Supplementary Table 1**). The top 200 ranked probes (p<0.05 and largest effect sizes) are listed in **Supplementary Table 2**.

A further 31,770 probes met criteria for longitudinal change unique to remitted ADHD (**Supplementary Figure 2a**), of which, 67% (n=21,213 probes) mapped to a single gene, again enriched in brain tissue (**Supplementary Figure 2b**), and 25 brain regions spanning prenatal through to adult life (**Supplementary Table 1**). The top 200 probes are summarized in **Supplementary Table 3**.

### Longitudinal change in DNA methylation between ADHD and controls

Using probes meeting criteria in the total cohort, uncorrected tests identified 14,469 probes with a main diagnostic effect, and 20,522 with a diagnosis interaction with time. However, these findings did not survive correction for multiple comparisons (**Supplementary Table 4)**.

When stratifying by diagnostic outcome, persistent ADHD showed significant hypomethylation relative to controls at two probes (cg15550572 – *PRKCZ*, cg12577151 – *ACAD9*), and time-dependent differences at three probes (cg26901352 – *CCL17*, cg17657037 – ITGB1BP3, and cg19892195 – *AGAP1*; **Figure 3**). Remitted ADHD differed significantly from controls in the average DNA methylation of three probes (cg11124426, cg02208776 – *HSPE1:HSPD1*, and cg12881363 – *MOV10L1*; **Table 2**).

**Figure 3.**
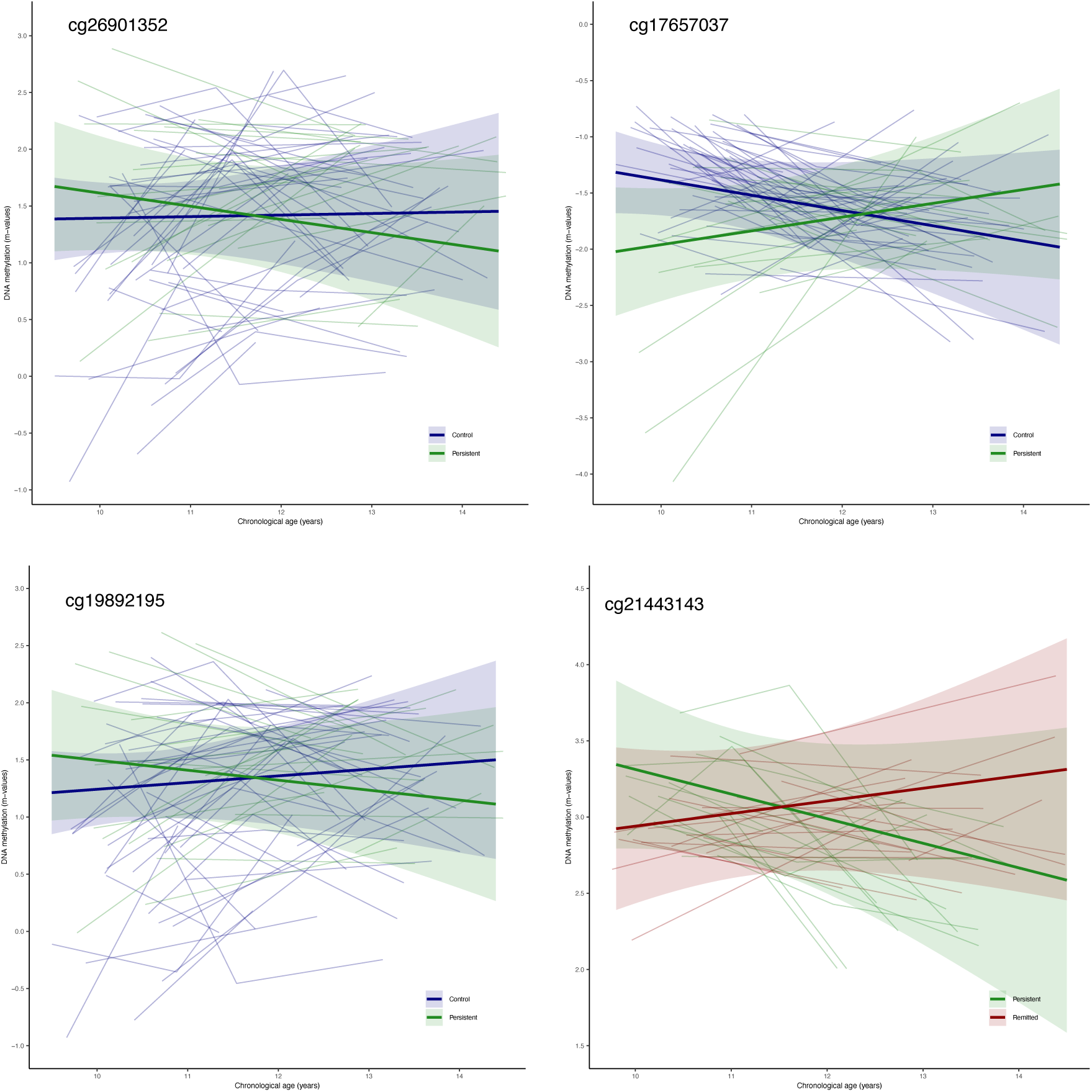
Longitudinal DNA methylation trajectories at CpG sites surviving FDR correction. Plots show methylation change over time for controls (blue), remitted ADHD (red) and persistent ADHD (green). Sites include cg26901352 – *CCL17*, cg17657037 – *ITGB1BP3*, and cg19892195 – *AGAP1*, with significant diagnosis x time interactions, and cg21443143 – *ZFAT*, which showed greater decline in remitted ADHD than in persistent ADHD.

**Table 2.** Differentially methylated probes between diagnostic groups. CpG probes showing significant main effects or diagnosis x time interactions in comparison between remitted ADHD, persistent ADHD, and control groups. Chromosomal location (Chr), strand orientation, associated gene, and effect sizes (logFC) with FDR-adjusted p-values provided.

| Comparisons | Probe ID | Chr | Position | Gene | Main<br>logFC<br>(p-value) | Interaction<br>logFC<br>(p-value) |
| --- | --- | --- | --- | --- | --- | --- |
| **Remitted vs.**<br>**controls** | cg11124426 | 2 | 131149108 |  | -0.67 (0.02) |  |
|  | cg02208776 | 2 | 198365119 | *HSPE1;HSPD1* | 0.83 (0.03) |  |
|  | cg12881363 | 22 | 50528298 | *MOV10L1* | 1.17 (0.03) |  |
| **Persistent vs.**<br>**controls** | cg15550572 | 1 | 2064441 | *PRKCZ* | -0.75 (0.05) |  |
|  | cg12577151 | 3 | 128615097 | *ACAD9* | -0.42 (0.05) |  |
|  | cg26901352 | 16 | 57447502 | *CCL17* |  | -0.13 (0.02) |
|  | cg17657037 | 19 | 3933856 | *ITGB1BP3* |  | 0.26 (0.03) |
|  | cg19892195 | 2 | 236963462 | *AGAP1* |  | -0.15 (0.04) |
| **Remitted vs.**<br>**persistent** | cg17601216 | 8 | 133787690 | *PHF20L1* | -0.58 (0.03) |  |
|  | cg00060320 | 3 | 134369974 | *KY* | -0.62 (0.02) |  |
|  | cg02968116 | 2 | 66652878 |  | 0.55 (0.03) |  |
|  | cg21443143 | 8 | 135491240 | *ZFAT* |  | -0.24 (0.02) |

### Change in DNA methylation unique to remitted ADHD

Among probes uniquely changing in remitted ADHD, three sites (cg17601216 – *PHF20L1*, cg00060320 – *KY*, cg02968116) showed significant main effects between remitted and persistent ADHD (**Table 2**). A fourth probe, cg21443143 – *ZFAT*, exhibited **greater** methylation decrease over time in remitted ADHD than in persistent ADHD (**Figure 3**). These genes are expressed broadly in cortical and subcortical brain regions, across developmental stages, with *PHF20L1* highly represented in neocortical areas (i.e., dorsolateral temporal neocortex, frontal neocortex, neocortex [isocortex] and temporal neocortex).

### Change in DNA methylation and change in ADHD symptom severity

To examine links between epigenetic and clinical change, we performed residual-change analyses on four significant diagnostic-outcome probes (cg17601216, cg00060320, cg02968116 and cg02968116). As expected, ADHD symptoms decreased more in remitted ADHD (-1.37 ± 4.40) than persistent ADHD (1.23 ± 2.92, *t* = -4.25, *p*<0.0001). However, no significant associations were observed between residual change in methylation and symptom change for three probes (cg17601216: *ß*=0.003 [95% CI: -0.03, 0.03], *p*=0.85; cg00060320: *ß*=-0.01 [-0.03, 0.02], *p*=0.46; cg02968116: *ß*=-0.01 [-0.03, 0.01], *p*=0.20). A positive trend was noted for cg21443143 - *ZFAT* (*ß*=0.032 [-0.01, 0.07], *p*=0.085).

### Gene Enrichment

Functional understanding of the top 200 probes (main diagnostic effect and diagnosis interaction with time) revealed 283 GO terms and two KEGG pathways, at a nominal p<0.05. These cover processes such as cerebellar maturation and neurotransmitter receptor transport – but none of these findings survived multiple correction (**Supplementary Table 5**).

For the diagnostic outcome findings, all 12 FDR-significant probes were analyzed for GO and KEGG enrichment (**Figure 4a and b**). Two probes annotated to *PRKCZ* and *AGAP1*, linked to long term memory (*PRKCZ*) or synaptic structure modification (*AGAP1*). *PRKCZ* was also enriched in the top-ranking chemokine signaling pathway, and associated with several disease classes, including mental disorders and nervous system diseases (**Figure 4c**), along with *HSPD1*, *ACAD9*, *ZFAT* and *MOV10L1*. As above, no enrichment results survived correction for multiple comparisons (**Supplementary Table 6**).

**Figure 4.**
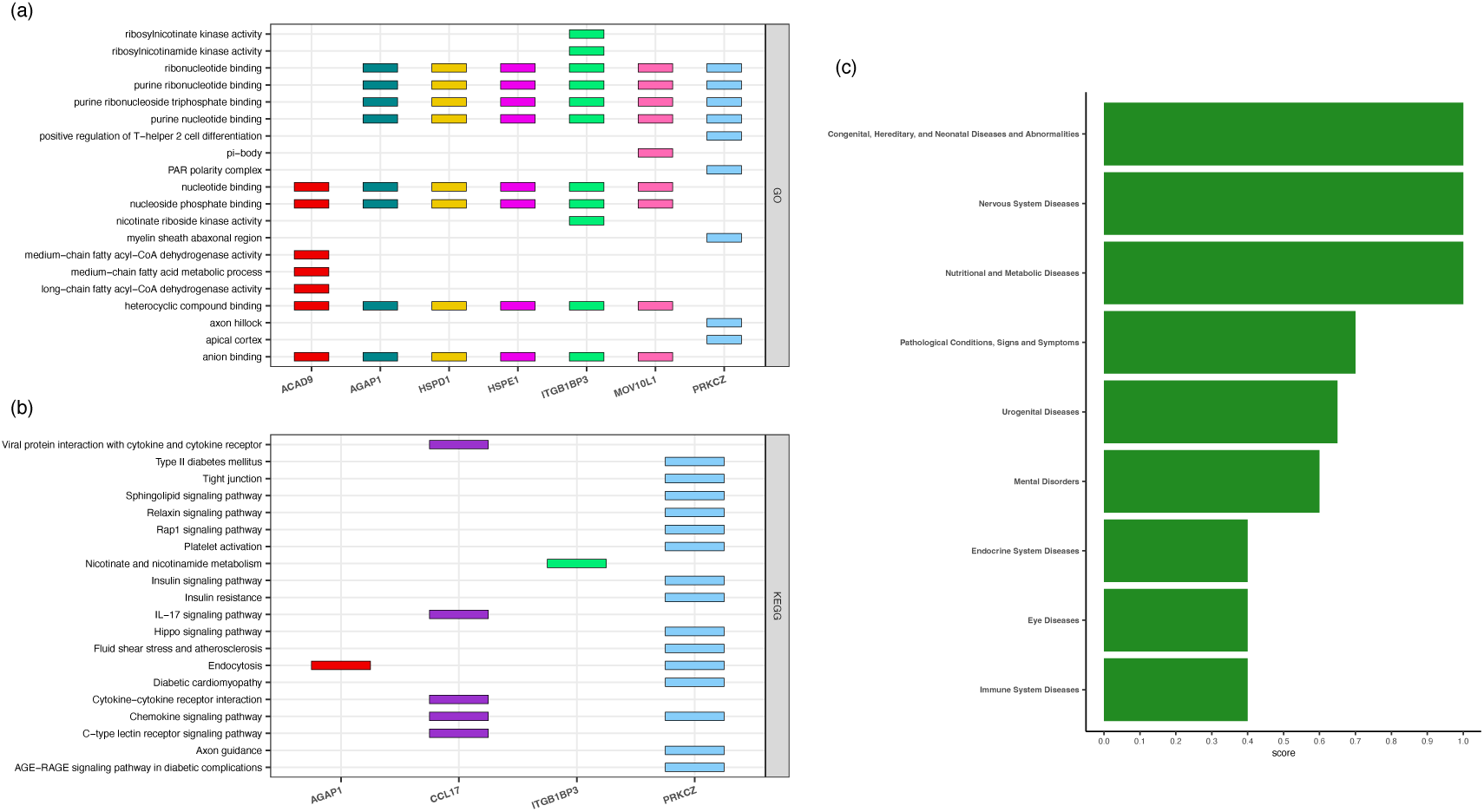
Functional enrichment analyses of differentially methylated CpGs. (a-b) Gene Ontology (GO) terms and KEGG pathways most enriched (nominal p<0.05) among FDR-significant probes. (C) Disease-association analysis linking probe-annotated genes (e.g., *PRKCZ*, *AGAP1*, *HSPD1*, *ACAD9*, *ZFAT*, and *MOV10L1*) to nervous system and mental disorders.

## DISCUSSION

This study examined whether longitudinal change in DNA methylation differs between children and adolescents with and without ADHD, and is among the first to explore the epigenetic patterns distinguishing persistent from remitted ADHD. By quantifying methylation across three developmental waves, we found no robust differences between ADHD cases and controls after multiple comparison correction, but we did identify 12 CpG sites that distinguished diagnostic subgroups from each other. Many of these probes map to genes expressed in brain tissue, suggesting biologically relevant methylation changes that may help explain variability in ADHD outcomes.

Most earlier prospective studies have focused on whether early life DNA methylation (measured from cord or whole blood) predicts late childhood symptoms (13, 18, 19). From these, two studies found an association with childhood ADHD symptoms when DNA methylation was measured at birth, but not when samples were collected concurrently during childhood (18, 19). Our null finding between a pooled ADHD sample and controls align with several saliva-based epigenome wide case-control or cross-sectional studies, which had also adjusted for multiple comparisons (17, 46, 47).

Symptoms of ADHD in childhood can vary over the course of adolescence, and individuals can remit or persist in their diagnostic status over time. To further understand this diversity at the molecular level, we compared DNA methylation between children and adolescents with remitted or persistent ADHD, and identified four differentially methylated sites. Among these, cg21443143 (annotated to *ZFAT*) showed a unique decline in methylation over time among remitted participants. *ZFAT* encodes a transcriptional regulator involved in immune cell survival and apoptosis (48), and autoimmune thyroid disease (49, 50). This aligns with evidence implicating immune processes and inflammation in ADHD pathophysiology (51–53). In relation to the brain, we found this gene to be enriched in several brain regions, most notable in the cingulate cortex and the prefrontal region, which echoes neuroimaging findings (54–58). This CpG site, however, was not identified by the only other published epigenetic study in remitted and persistent ADHD, though this was a case-control study involving a relatively small sample of young adults (21). Therefore, our finding warrants further investigation using repeated measures in children and adolescents.

In comparison to controls, a most consistent finding for persistent ADHD was a lower or declining methylation over time. Three of the identified sites map to genes enriched in brain tissue collected during infancy and childhood (*PRKCZ, ACAD9 and AGAP1*), including the somatosensory cortex, which supports prior NICAP neuroimaging data (59). Functionally, some genes are implicated in synaptic structure modification (*AGAP1*) and long-term memory (*PRKCZ*), which are essential for brain development and can be impaired in persons with ADHD (60, 61). Whether the lower DNA methylation of these genes represents a heightened expression that ultimately maintains symptom severity in ADHD is yet to be seen, though some evidence has shown overexpression of *AGAP1* to have a negative impact on dendritic spine density (62).

CpG sites distinguishing remitted ADHD from controls were mostly hypermethylated, though none of these maps to brain-expressed genes. One site near the *HSPD1* gene (cg02208776), however, has been previously associated with brain morphology in schizophrenia (63) and may have neurodevelopmental relevance. The scarcity of research limits our interpretation of this finding, and it should be noted that gene expression was not directly measured in the present study, and therefore these sites in remitted ADHD could still be linked directly to the brain with further studies. In the previously mentioned study by Meijer et al (2020), which involved small samples of cases and controls, they found no evidence of a difference between individuals with remitted ADHD and controls, though investigations using large cohort studies, and therefore with greater power would provide further insight.

The current study is one of the few to investigate longitudinal change in DNA methylation across the epigenome, and compare these changes between children and adolescents with or without ADHD. It is also the first to explore DNA methylation in relation to change in diagnostic outcomes (i.e., persistence and remitted). However, while this cohort had data across three waves, most individuals provided only two saliva samples, and across a limited age range. Moreover, although these findings are limited to epigenetic markers measured from saliva, and therefore cannot inform directly on DNA methylation in brain tissue, studies do report relative stability in DNA methylation across tissue types (64). We address this gap by testing the enrichment of genes within brain regions of the Allan Brain Atlas, which defines gene expression at varying stages across the lifespan (35). Further studies on these sites are planned to explore links to the brain by leveraging the extensive neuroimaging data collected through the NICAP study.

## CONCLUSION

Our study identified several epigenetic differences between children and adolescents with persistent or remitted ADHD, from neurotypical peers, and from each other. Given the novelty of these findings, future research may benefit from epigenetic investigations in diagnostic subgroups, rather than pooled clinical samples, using data collective at repeated timepoints.

## ACKNOWLEDGEMENTS

The work was funded by the National Health and Medical Research Council of Australia (NHMRC; grant #2029361) and a grant from the Waterloo Foundation.

The broader Children’s Attention Project (CAP) and the Neuroimaging of the Children’s Attention Project (NICAP) cohort was funded by NHMRC (grants #1065895 and #1008522).

A special thanks to all the children and families for their participation in this study. This work was supported by the MASSIVE HPC facility (www.massive.org.au).

## CONFLICT OF INTEREST

The authors declare no conflict of interest.

## Notes

### Competing Interest Statement

The authors have declared no competing interest.

## REFERENCES

1. Cortese S, Song M, Farhat LC, Yon DK, Lee SW, Kim MS, et al. Incidence, prevalence, and global burden of ADHD from 1990 to 2019 across 204 countries: data, with critical re-analysis, from the Global Burden of Disease study. Mol Psychiatry. 2023;28(11):4823–30.

2. Sibley MH, Arnold LE, Swanson JM, Hechtman LT, Kennedy TM, Owens E, et al. Variable Patterns of Remission From ADHD in the Multimodal Treatment Study of ADHD. Am J Psychiatry. 2022;179(2):142–51.

3. Fayyad J, Sampson NA, Hwang I, Adamowski T, Aguilar-Gaxiola S, Al-Hamzawi A, et al. The descriptive epidemiology of DSM-IV Adult ADHD in the World Health Organization World Mental Health Surveys. Atten Defic Hyperact Disord. 2017;9(1):47–65.

4. Kuntsi J, Rijsdijk F, Ronald A, Asherson P, Plomin R. Genetic influences on the stability of attention-deficit/hyperactivity disorder symptoms from early to middle childhood. Biol Psychiatry. 2005;57(6):647–54.

5. Riglin L, Collishaw S, Thapar AK, Dalsgaard S, Langley K, Smith GD, et al. Association of Genetic Risk Variants With Attention-Deficit/Hyperactivity Disorder Trajectories in the General Population. JAMA Psychiatry. 2016;73(12):1285–92.

6. Roy A, Hechtman L, Arnold LE, Sibley MH, Molina BS, Swanson JM, et al. Childhood Factors Affecting Persistence and Desistence of Attention-Deficit/Hyperactivity Disorder Symptoms in Adulthood: Results From the MTA. J Am Acad Child Adolesc Psychiatry. 2016;55(11):937–44 e4.

7. Cecil CAM, Nigg JT. Epigenetics and ADHD: Reflections on Current Knowledge, Research Priorities and Translational Potential. Mol Diagn Ther. 2022;26(6):581–606.

8. Walton E, Baltramonaityte V, Calhoun V, Heijmans BT, Thompson PM, Cecil CAM. A systematic review of neuroimaging epigenetic research: calling for an increased focus on development. Mol Psychiatry. 2023;28(7):2839–47.

9. Wheater ENW, Stoye DQ, Cox SR, Wardlaw JM, Drake AJ, Bastin ME, et al. DNA methylation and brain structure and function across the life course: A systematic review. Neurosci Biobehav Rev. 2020;113:133–56.

10. Carvalho G, Costa T, Nascimento AM, Wolff BM, Damasceno JG, Vieira LL, et al. DNA methylation epi-signature and biological age in attention deficit hyperactivity disorder patients. Clin Neurol Neurosurg. 2023;228:107714.

11. Chen YC, Sudre G, Sharp W, Donovan F, Chandrasekharappa SC, Hansen N, et al. Neuroanatomic, epigenetic and genetic differences in monozygotic twins discordant for attention deficit hyperactivity disorder. Mol Psychiatry. 2018;23(3):683–90.

12. Ehlinger JV, Goodrich JM, Dolinoy DC, Watkins DJ, Cantoral A, Mercado-Garcia A, et al. Associations between blood leukocyte DNA methylation and sustained attention in mid-to-late childhood. Epigenomics. 2023;15(19):965–81.

13. Camerota M, Lester BM, Castellanos FX, Carter BS, Check J, Helderman J, et al. Epigenome-wide association study identifies neonatal DNA methylation associated with two-year attention problems in children born very preterm. Transl Psychiatry. 2024;14(1):126.

14. Gervin K, Nordeng H, Ystrom E, Reichborn-Kjennerud T, Lyle R. Long-term prenatal exposure to paracetamol is associated with DNA methylation differences in children diagnosed with ADHD. Clin Epigenetics. 2017;9:77.

15. Goodman SJ, Burton CL, Butcher DT, Siu MT, Lemire M, Chater-Diehl E, et al. Obsessive-compulsive disorder and attention-deficit/hyperactivity disorder: distinct associations with DNA methylation and genetic variation. Journal of neurodevelopmental disorders. 2020;12:1–15.

16. Li SC, Kuo HC, Huang LH, Chou WJ, Lee SY, Chan WC, et al. DNA Methylation in LIME1 and SPTBN2 Genes Is Associated with Attention Deficit in Children. Children (Basel). 2021;8(2).

17. Mooney MA, Ryabinin P, Wilmot B, Bhatt P, Mill J, Nigg JT. Large epigenome-wide association study of childhood ADHD identifies peripheral DNA methylation associated with disease and polygenic risk burden. Transl Psychiatry. 2020;10(1):8.

18. Neumann A, Walton E, Alemany S, Cecil C, Gonzalez JR, Jima DD, et al. Association between DNA methylation and ADHD symptoms from birth to school age: a prospective meta-analysis. Transl Psychiatry. 2020;10(1):398.

19. Walton E, Pingault JB, Cecil CA, Gaunt TR, Relton CL, Mill J, et al. Epigenetic profiling of ADHD symptoms trajectories: a prospective, methylome-wide study. Mol Psychiatry. 2017;22(2):250–6.

20. Wilmot B, Fry R, Smeester L, Musser ED, Mill J, Nigg JT. Methylomic analysis of salivary DNA in childhood ADHD identifies altered DNA methylation in VIPR2. J Child Psychol Psychiatry. 2016;57(2):152–60.

21. Meijer M, Klein M, Hannon E, Van der Meer D, Hartman C, Oosterlaan J, et al. Genome-wide DNA methylation patterns in persistent attention-deficit/hyperactivity disorder and in association with impulsive and callous traits. Frontiers in genetics. 2020;11:16.

22. Silk TJ, Genc S, Anderson V, Efron D, Hazell P, Nicholson JM, et al. Developmental brain trajectories in children with ADHD and controls: a longitudinal neuroimaging study. BMC Psychiatry. 2016;16:59.

23. Sciberras E, Efron D, Schilpzand EJ, Anderson V, Jongeling B, Hazell P, et al. The Children’s Attention Project: a community-based longitudinal study of children with ADHD and non-ADHD controls. BMC Psychiatry. 2013;13:18.

24. Conners CK. Conners third edition (Conners 3). Los Angeles, CA: Western Psychological Services. 2008.

25. Shaffer D, Fisher P, Lucas CP, Dulcan MK, Schwab-Stone ME. NIMH Diagnostic Interview Schedule for Children Version IV (NIMH DISC-IV): description, differences from previous versions, and reliability of some common diagnoses. J Am Acad Child Adolesc Psychiatry. 2000;39(1):28–38.

26. Silk TJ, Genc S, Anderson V, Efron D, Hazell P, Nicholson JM, et al. Developmental brain trajectories in children with ADHD and controls: a longitudinal neuroimaging study. BMC Psychiatry. 2016;16:1–9.

27. Aryee MJ, Jaffe AE, Corrada-Bravo H, Ladd-Acosta C, Feinberg AP, Hansen KD, et al. Minfi: a flexible and comprehensive Bioconductor package for the analysis of Infinium DNA methylation microarrays. Bioinformatics. 2014;30(10):1363–9.

28. Touleimat NT, J. Complete pipeline for Infinium® Human Methylation 450K BeadChip data processing using subset quantile normalization for accurate DNA methylation estimation. Epigenomics. 2012;4(3):325–41.

29. Wu Z, Aryee MJ. Subset quantile normalization using negative control features. J Comput Biol. 2010;17(10):1385–95.

30. Pidsley R, Zotenko E, Peters TJ, Lawrence MG, Risbridger GP, Molloy P, et al. Critical evaluation of the Illumina MethylationEPIC BeadChip microarray for whole-genome DNA methylation profiling. Genome Biol. 2016;17(1):208.

31. Du P, Zhang X, Huang CC, Jafari N, Kibbe WA, Hou L, et al. Comparison of Beta-value and M-value methods for quantifying methylation levels by microarray analysis. BMC Bioinformatics. 2010;11:587.

32. Teschendorff A, Zheng S. Epigenetic dissection of intra-sample-heterogeneity (EPIDISH). 2023.

33. Uhlen M, Fagerberg L, Hallstrom BM, Lindskog C, Oksvold P, Mardinoglu A, et al. Proteomics. Tissue-based map of the human proteome. Science. 2015;347(6220):1260419.

34. Jain A, Tuteja G. TissueEnrich: Tissue-specific gene enrichment analysis. Bioinformatics. 2019;35(11):1966–7.

35. Grote S, Prufer K, Kelso J, Dannemann M. ABAEnrichment: an R package to test for gene set expression enrichment in the adult and developing human brain. Bioinformatics. 2016;32(20):3201–3.

36. Ashburner M, Ball CA, Blake JA, Botstein D, Butler H, Cherry JM, et al. Gene ontology: tool for the unification of biology. The Gene Ontology Consortium. Nat Genet. 2000;25(1):25–9.

37. Gene Ontology C, Aleksander SA, Balhoff J, Carbon S, Cherry JM, Drabkin HJ, et al. The Gene Ontology knowledgebase in 2023. Genetics. 2023;224(1).

38. Kanehisa M, Goto S. KEGG: kyoto encyclopedia of genes and genomes. Nucleic Acids Res. 2000;28(1):27–30.

39. Phipson B, Maksimovic J, Oshlack A. missMethyl: an R package for analyzing data from Illumina’s HumanMethylation450 platform. Bioinformatics. 2016;32(2):286–8.

40. Landrum MJ, Lee JM, Benson M, Brown GR, Chao C, Chitipiralla S, et al. ClinVar: improving access to variant interpretations and supporting evidence. Nucleic Acids Res. 2018;46(D1):D1062–D7.

41. Rehm HL, Berg JS, Brooks LD, Bustamante CD, Evans JP, Landrum MJ, et al. ClinGen--the Clinical Genome Resource. N Engl J Med. 2015;372(23):2235–42.

42. Gutierrez-Sacristan A, Bravo A, Portero-Tresserra M, Valverde O, Armario A, Blanco-Gandia MC, et al. Text mining and expert curation to develop a database on psychiatric diseases and their genes. Database (Oxford). 2017;2017.

43. Pinero J, Ramirez-Anguita JM, Sauch-Pitarch J, Ronzano F, Centeno E, Sanz F, et al. The DisGeNET knowledge platform for disease genomics: 2019 update. Nucleic Acids Res. 2020;48(D1):D845–D55.

44. Benjamini Y, Hochberg Y. Controlling the false discovery rate: a practical and powerful approach to multiple testing. Journal of the Royal statistical society: series B (Methodological). 1995;57(1):289–300.

45. R Core Team. R: A Language and Environment for Statistical Computing. Vienna, Austria: R Foundation for Statistical Computing; 2024.

46. Fujisawa TX, Nishitani S, Makita K, Yao A, Takiguchi S, Hamamura S, et al. Association of Epigenetic Differences Screened in a Few Cases of Monozygotic Twins Discordant for Attention-Deficit Hyperactivity Disorder With Brain Structures. Front Neurosci. 2021;15:799761.

47. Goodman SJ, Burton CL, Butcher DT, Siu MT, Lemire M, Chater-Diehl E, et al. Obsessive-compulsive disorder and attention-deficit/hyperactivity disorder: distinct associations with DNA methylation and genetic variation. J Neurodev Disord. 2020;12(1):23.

48. Fujimoto T, Doi K, Koyanagi M, Tsunoda T, Takashima Y, Yoshida Y, et al. ZFAT is an antiapoptotic molecule and critical for cell survival in MOLT-4 cells. FEBS Lett. 2009;583(3):568–72.

49. Koyanagi M, Nakabayashi K, Fujimoto T, Gu N, Baba I, Takashima Y, et al. ZFAT expression in B and T lymphocytes and identification of ZFAT-regulated genes. Genomics. 2008;91(5):451–7.

50. Shirasawa S, Harada H, Furugaki K, Akamizu T, Ishikawa N, Ito K, et al. SNPs in the promoter of a B cell-specific antisense transcript, SAS-ZFAT, determine susceptibility to autoimmune thyroid disease. Hum Mol Genet. 2004;13(19):2221–31.

51. Han VX, Patel S, Jones HF, Nielsen TC, Mohammad SS, Hofer MJ, et al. Maternal acute and chronic inflammation in pregnancy is associated with common neurodevelopmental disorders: a systematic review. Transl Psychiatry. 2021;11(1):71.

52. Zhong Y, Baum LW, Tubbs JD, Ye R, Chen LH, Wu T, et al. Common and rare variant analyses implicate late-infancy cerebellar development and immune genes in ADHD. J Neurodev Disord. 2025;17(1):34.

53. Liu Z, Wang L, Yu L, Zhao Y, Zhu M, Wang Y, et al. Identification of immune cells and circulating inflammatory factors associated with neurodevelopmental disorders by bidirectional Mendelian randomization and mediation analysis. Sci Rep. 2025;15(1):12840.

54. Clerkin SM, Schulz KP, Berwid OG, Fan J, Newcorn JH, Tang CY, et al. Thalamo-cortical activation and connectivity during response preparation in adults with persistent and remitted ADHD. Am J Psychiatry. 2013;170(9):1011–9.

55. Mattfeld AT, Gabrieli JD, Biederman J, Spencer T, Brown A, Kotte A, et al. Brain differences between persistent and remitted attention deficit hyperactivity disorder. Brain. 2014;137(Pt 9):2423–8.

56. Schulz KP, Li X, Clerkin SM, Fan J, Berwid OG, Newcorn JH, et al. Prefrontal and parietal correlates of cognitive control related to the adult outcome of attention-deficit/hyperactivity disorder diagnosed in childhood. Cortex. 2017;90:1–11.

57. Shaw P, Malek M, Watson B, Greenstein D, de Rossi P, Sharp W. Trajectories of cerebral cortical development in childhood and adolescence and adult attention-deficit/hyperactivity disorder. Biol Psychiatry. 2013;74(8):599–606.

58. Wetterling F, McCarthy H, Tozzi L, Skokauskas N, O’Doherty JP, Mulligan A, et al. Impaired reward processing in the human prefrontal cortex distinguishes between persistent and remittent attention deficit hyperactivity disorder. Hum Brain Mapp. 2015;36(11):4648–63.

59. Fuelscher I, Hyde C, Thomson P, Vijayakumar N, Sciberras E, Efron D, et al. Longitudinal Trajectories of White Matter Development in Attention-Deficit/Hyperactivity Disorder. Biol Psychiatry Cogn Neurosci Neuroimaging. 2023;8(11):1103–12.

60. Mohammadkhanloo M, Pooyan M, Sharini H, Yousefpour M. Investigating resting-state functional connectivity changes within procedural memory network across neuropsychiatric disorders using fMRI. BMC Med Imaging. 2025;25(1):18.

61. Rhodes SM, Park J, Seth S, Coghill DR. A comprehensive investigation of memory impairment in attention deficit hyperactivity disorder and oppositional defiant disorder. J Child Psychol Psychiatry. 2012;53(2):128–37.

62. Arnold M, Cross R, Singleton KS, Zlatic S, Chapleau C, Mullin AP, et al. The Endosome Localized Arf-GAP AGAP1 Modulates Dendritic Spine Morphology Downstream of the Neurodevelopmental Disorder Factor Dysbindin. Front Cell Neurosci. 2016;10:218.

63. van der Meer D, Shadrin AA, O’Connell K, Bettella F, Djurovic S, Wolfers T, et al. Boosting Schizophrenia Genetics by Utilizing Genetic Overlap With Brain Morphology. Biol Psychiatry. 2022;92(4):291–8.

64. Nishitani S, Isozaki M, Yao A, Higashino Y, Yamauchi T, Kidoguchi M, et al. Cross-tissue correlations of genome-wide DNA methylation in Japanese live human brain and blood, saliva, and buccal epithelial tissues. Transl Psychiatry. 2023;13(1):72.

