## Supplementary Figures 1-2 and Supplementary Tables 1-6 for "Longitudinal DNA methylation dynamics distinguish Persistent and Remitted ADHD in childhood and adolescence"

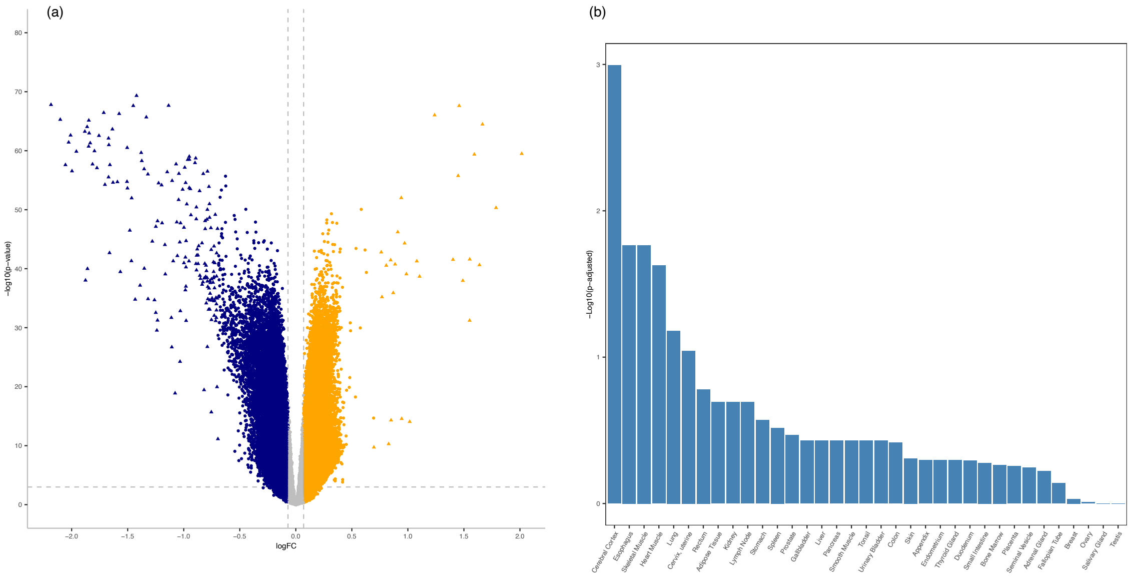


**Supplementary Figure 1. DNA methylation profiles for CpGs with significant change across development.** Volcano plot (a) and bar plot for tissue enrichment results (b) for probes meeting criteria for change ($p<0.05$ and >5% change) in full study sample.

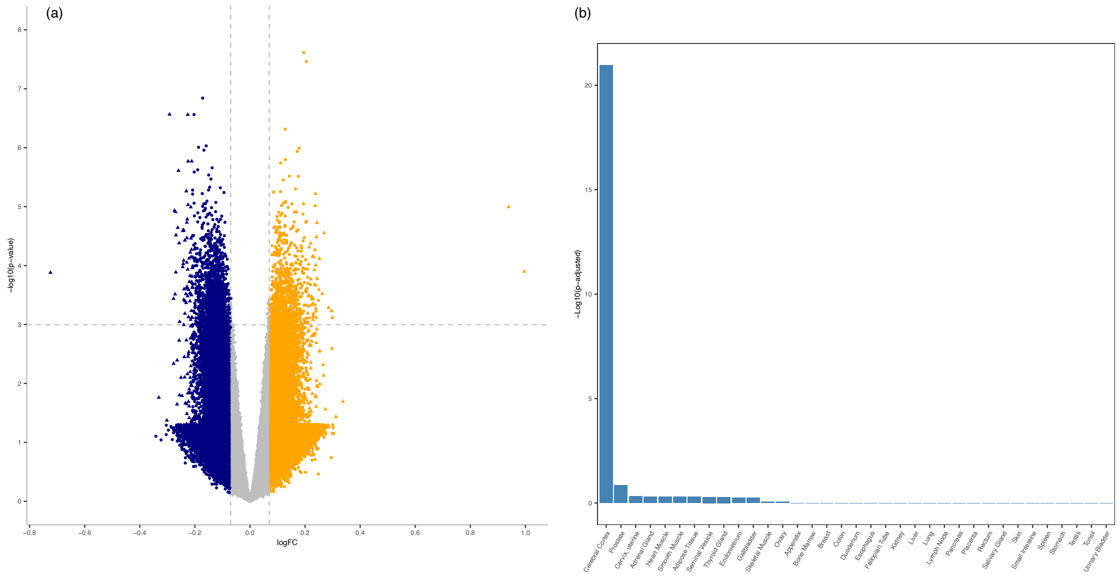


**Supplementary Figure 2. DNA methylation profiles for CpGs with significant change unique to remitted ADHD.** Volcano plot (a) and bar plot for tissue enrichment results (b) for probes meeting criteria for change ($p<0.05$ and >5% change) unique to remitted ADHD.

### SUPPLEMENTARY TABLES

**Supplementary Table 1. Gene enrichment in brain tissue collected across the lifespan.** Enrichment in brain regions of the Allan Brain Atlas (Brain Structure ID and Brain Structure) consists of genes annotated to probes that met our study criteria for longitudinal change in the total sample (*'Longitudinal change CpGs'*) or was unique to remitted ADHD (*'Diagnostic outcomes CpGs'*). Results limited to min False-Wise Error Rate <0.05.

| Age category | Brain Structure ID | Brain Structure | min FWER | Probes |
|---|---|---|---|---|
| *Longitudinal change CpGs* | | | | |
| 0-2yrs | Allen:10269 | Primary visual cortex (striate cortex, area V1/17) | <0.00001 | 91672 |
| 0-2yrs | Allen:10331 | Cerebral nuclei | 0.002 | 43490 |
| 0-2yrs | Allen:10163 | Primary motor cortex (area M1, area 4) | 0.004 | 95795 |
| 0-2yrs | Allen:10185 | Ventrolateral prefrontal cortex | 0.01 | 95091 |
| 0-2yrs | Allen:10208 | Parietal neocortex | 0.02 | 92400 |
| 0-2yrs | Allen:10209 | Primary somatosensory cortex (area S1, areas 3,1,2) | 0.03 | 91009 |
| 3-11yrs | Allen:10657 | Cerebellar cortex | 0.01 | 76979 |
| *Diagnostic outcomes CpGs* | | | | |
| Prenatal | Allen:10252 | Inferolateral temporal cortex (area TEv, area 20) | 0.002 | 15924 |
| Prenatal | Allen:10243 | Posterior (caudal) superior temporal cortex (area 22c) | 0.01 | 14221 |
| Prenatal | Allen:10160 | Neocortex (isocortex) | 0.02 | 13172 |
| Prenatal | Allen:10225 | Posteroventral (inferior) parietal cortex | 0.02 | 15664 |
| Prenatal | Allen:10161 | Frontal neocortex | 0.03 | 10860 |
| Prenatal | Allen:10163 | Primary motor cortex (area M1, area 4) | 0.03 | 14089 |
| Prenatal | Allen:10278 | Anterior (rostral) cingulate (medial prefrontal) cortex | 0.03 | 12571 |
| Prenatal | Allen:10235 | Temporal neocortex | 0.04 | 14497 |
| Prenatal | Allen:10158 | Telencephalon | 0.05 | 7941 |
| Prenatal | Allen:10333 | Striatum | 0.05 | 16202 |
| Prenatal | Allen:10194 | Orbital frontal cortex | 0.05 | 15991 |
| Prenatal | Allen:10331 | Cerebral nuclei | 0.05 | 16345 |
| 0-2yrs | Allen:10331 | Cerebral nuclei | 0.001 | 15220 |
| 0-2yrs | Allen:10185 | Ventrolateral prefrontal cortex | 0.001 | 16121 |

| | | | | |
|---|---|---|---|---|
| 0-2yrs | Allen:10243 | Posterior (caudal) superior temporal cortex (area 22c) | 0.001 | 16055 |
| 0-2yrs | Allen:10194 | Orbital frontal cortex | 0.001 | 14447 |
| 0-2yrs | Allen:10269 | Primary visual cortex (striate cortex, area V1/17) | 0.003 | 10261 |
| 0-2yrs | Allen:10209 | Primary somatosensory cortex (area S1, areas 3,1,2) | 0.004 | 14331 |
| 0-2yrs | Allen:10278 | Anterior (rostral) cingulate (medial prefrontal) cortex | 0.01 | 15975 |
| 0-2yrs | Allen:10236 | Primary auditory cortex (core) | 0.01 | 14641 |
| 0-2yrs | Allen:10294 | Hippocampus (hippocampal formation) | 0.01 | 16273 |
| 0-2yrs | Allen:10225 | Posteroventral (inferior) parietal cortex | 0.02 | 15966 |
| 0-2yrs | Allen:10172 | Prefrontal cortex | 0.02 | 14706 |
| 0-2yrs | Allen:10157 | Gray matter of forebrain | 0.02 | 12318 |
| 0-2yrs | Allen:10208 | Parietal neocortex | 0.03 | 14488 |
| 0-2yrs | Allen:13322 | Dorsolateral temporal neocortex | 0.03 | 14701 |
| 0-2yrs | Allen:10252 | Inferolateral temporal cortex (area TEv, area 20) | 0.04 | 16042 |
| 0-2yrs | Allen:10163 | Primary motor cortex (area M1, area 4) | <0.00001 | 16154 |
| 0-2yrs | Allen:10361 | Amygdaloid complex | <0.00001 | 16350 |
| 0-2yrs | Allen:10398 | Mediodorsal nucleus of thalamus | <0.00001 | 15917 |
| 3-11yrs | Allen:10194 | Orbital frontal cortex | 0.002 | 12639 |
| 3-11yrs | Allen:10236 | Primary auditory cortex (core) | 0.01 | 10393 |
| 3-11yrs | Allen:10235 | Temporal neocortex | 0.02 | 12874 |
| 3-11yrs | Allen:10278 | Anterior (rostral) cingulate (medial prefrontal) cortex | 0.03 | 7591 |
| 3-11yrs | Allen:10157 | Gray matter of forebrain | 0.04 | 11977 |
| 3-11yrs | Allen:10269 | Primary visual cortex (striate cortex, area V1/17) | <0.00001 | 12587 |
| 3-11yrs | Allen:10657 | Cerebellar cortex | <0.00001 | 8576 |
| 12-19yrs | Allen:10173 | Dorsolateral prefrontal cortex | 0.004 | 16023 |
| 12-19yrs | Allen:10236 | Primary auditory cortex (core) | 0.004 | 16035 |
| 12-19yrs | Allen:10294 | Hippocampus (hippocampal formation) | 0.01 | 16180 |
| 12-19yrs | Allen:10278 | Anterior (rostral) cingulate (medial prefrontal) cortex | 0.01 | 16051 |
| 12-19yrs | Allen:10361 | Amygdaloid complex | 0.01 | 16147 |
| 12-19yrs | Allen:10194 | Orbital frontal cortex | 0.04 | 16019 |
| >19yrs | Allen:10225 | Posteroventral (inferior) parietal cortex | 0.01 | 12485 |
| >19yrs | Allen:10185 | Ventrolateral prefrontal cortex | 0.02 | 12586 |
| >19yrs | Allen:10278 | Anterior (rostral) cingulate (medial prefrontal) cortex | 0.03 | 12635 |
| >19yrs | Allen:10243 | Posterior (caudal) superior temporal cortex (area 22c) | 0.04 | 12535 |

**Supplementary Table 2. Top 200 ranked probes meeting criteria for longitudinal change (p<0.05 and >5%) in the full study sample.**

Chromosomal location (Chr), strand orientation, associated gene, and effect sizes (logFC) with unadjusted p-values provided.

| Probes | Chr | Position | *Gene* | logFC | p-value |
|---|---|---|---|---|---|
| cg08452456 | chr17 | 45786439 | *TBKBP1* | -2.18 | <0.00001 |
| cg01483824 | chr19 | 48945886 | *GRIN2D* | -2.10 | <0.00001 |
| cg07538239 | chr11 | 1467081 | *BRSK2* | -2.06 | <0.00001 |
| cg00007036 | chr1 | 33742089 | *ZNF362* | -2.03 | <0.00001 |
| cg00160667 | chr1 | 29587357 | *PTPRU* | 2.02 | <0.00001 |
| cg19541688 | chr19 | 46289559 | *DMWD* | -2.01 | <0.00001 |
| cg05166248 | chr11 | 134252845 | *B3GAT1* | -2.00 | <0.00001 |
| cg07277190 | chr12 | 48723631 | *H1FNT* | -1.96 | <0.00001 |
| cg23367339 | chr17 | 36622717 | *ARHGAP23* | -1.88 | <0.00001 |
| cg21004358 | chr4 | 187422119 | | -1.88 | <0.00001 |
| cg16034787 | chr5 | 508349 | *SLC9A3* | -1.86 | <0.00001 |
| cg19081270 | chr7 | 4830343 | *KIAA0415* | -1.86 | <0.00001 |
| cg24534173 | chr17 | 3375043 | *SPATA22* | -1.85 | <0.00001 |
| cg13438128 | chr21 | 46293863 | *PTTG1IP* | -1.85 | <0.00001 |
| cg14859324 | chr15 | 26874363 | *GABRB3* | -1.84 | <0.00001 |
| cg23714330 | chr17 | 73084871 | *SLC16A5* | -1.83 | <0.00001 |
| cg15837531 | chr17 | 9162271 | *STX8* | -1.81 | <0.00001 |
| cg15866800 | chr6 | 147525459 | *STXBP5;STXBP5-AS1* | -1.80 | <0.00001 |
| cg22846334 | chr15 | 43941042 | *CATSPER2* | 1.79 | <0.00001 |
| cg05174710 | chr21 | 36164748 | *RUNX1* | -1.77 | <0.00001 |
| cg01666550 | chr11 | 68183850 | *LRP5* | -1.76 | <0.00001 |
| cg24954895 | chr5 | 140800761 | *PCDHGA1;PCDHGB1;PCDHGA2;PCDHGB2;PCDHGA3;PCDHGB3;PCDHGA4;PCDHGB4;PCDHGA5;PCDHGB5;PCDHGA6;PCDHGB6;PCDHGA7;PCDHGB7;PCDHGA8;PCDHGA9;PCDHGA10;PCDHGA11;* | -1.71 | <0.00001 |

| | | | | | |
|---|---|---|---|---|---|
| cg01504215 | chr2 | 238395781 | *MLPH* | -1.70 | <0.00001 |
| cg23841819 | chr1 | 204970383 | *NFASC* | -1.67 | <0.00001 |
| cg09706833 | chr21 | 46824938 | *COL18A1* | -1.67 | <0.00001 |
| cg14080585 | chr20 | 60639721 | *TAF4* | -1.67 | <0.00001 |
| cg09607566 | chr16 | 89367323 | *ANKRD11* | 1.67 | <0.00001 |
| cg04608651 | chr17 | 40557508 | *PTRF* | -1.66 | <0.00001 |
| cg09282336 | chr16 | 81526954 | *CMIP* | -1.66 | <0.00001 |
| cg03338348 | chr16 | 67840472 | *TSNAXIP1;RANBP10* | 1.64 | <0.00001 |
| cg05262724 | chr5 | 31908436 | *PDZD2* | -1.64 | <0.00001 |
| cg20227471 | chr2 | 25065550 | *ADCY3* | -1.63 | <0.00001 |
| cg13786191 | chr5 | 176057738 | *MIR4281;SNCB;EIF4E1B* | 1.59 | <0.00001 |
| cg01392253 | chr14 | 67708096 | *MPP5* | -1.59 | <0.00001 |
| cg01878345 | chr2 | 105471960 | *POU3F3* | -1.58 | <0.00001 |
| cg02613380 | chr10 | 99330076 | *UBTD1* | -1.57 | <0.00001 |
| cg09698465 | chr12 | 133000178 | | 1.55 | <0.00001 |
| cg05186755 | chr6 | 33217183 | | 1.55 | <0.00001 |
| cg01311063 | chr2 | 131058184 | | -1.51 | <0.00001 |
| cg11582634 | chr9 | 138594185 | *KCNT1* | -1.50 | <0.00001 |
| cg13981054 | chr6 | 18020115 | | -1.50 | <0.00001 |
| cg08689926 | chr22 | 50322930 | | 1.49 | <0.00001 |
| cg23879118 | chr11 | 67414373 | *ACY3* | -1.48 | <0.00001 |
| cg18710053 | chr5 | 126409061 | *FLJ44606* | -1.47 | <0.00001 |
| cg11843188 | chr11 | 102322558 | *TMEM123* | -1.46 | <0.00001 |
| cg09932386 | chr15 | 70433602 | | 1.46 | <0.00001 |
| cg04097963 | chr15 | 40616030 | | -1.45 | <0.00001 |
| cg04581613 | chr16 | 1543690 | *TELO2* | 1.45 | <0.00001 |
| cg26703758 | chr11 | 17791761 | *KCNC1* | -1.43 | <0.00001 |
| cg27443867 | chr10 | 43725376 | *RASGEF1A* | -1.42 | <0.00001 |
| cg27456487 | chr17 | 56349062 | *MPO* | 1.40 | <0.00001 |
| cg13777502 | chr17 | 66031814 | *KPNA2* | -1.39 | <0.00001 |
| cg11288801 | chr19 | 1406661 | *DAZAP1* | -1.38 | <0.00001 |
| cg02851212 | chr10 | 1451418 | *ADARB2* | -1.37 | <0.00001 |

| | | | | | |
|---|---|---|---|---|---|
| cg19529823 | chr2 | 10442953 | *HPCAL1* | -1.35 | <0.00001 |
| cg13464717 | chr19 | 2269532 | *OAZ1* | -1.35 | <0.00001 |
| cg27328757 | chr2 | 11295421 | *PQLC3* | -1.33 | <0.00001 |
| cg25485991 | chr17 | 8066461 | *VAMP2* | -1.32 | <0.00001 |
| cg24942922 | chr3 | 120003547 | | -1.32 | <0.00001 |
| cg23327483 | chr22 | 24110105 | *CHCHD10* | -1.28 | <0.00001 |
| cg22162544 | chr1 | 159842863 | *CFAP45* | -1.26 | <0.00001 |
| cg09629753 | chr16 | 29323721 | *RUNDC2C* | -1.25 | <0.00001 |
| cg12835012 | chr4 | 183795785 | | -1.25 | <0.00001 |
| cg03586128 | chr13 | 113299796 | | -1.24 | <0.00001 |
| cg02833127 | chr4 | 177116733 | *SPATA4* | 1.24 | <0.00001 |
| cg17682313 | chr4 | 153448438 | *FBXW7* | -1.23 | <0.00001 |
| cg24853765 | chr1 | 109584962 | *WDR47* | -1.23 | <0.00001 |
| cg15821589 | chr1 | 78444904 | *FUBP1* | -1.22 | <0.00001 |
| cg04534345 | chr4 | 82392459 | *RASGEF1B* | -1.20 | <0.00001 |
| cg07451781 | chr10 | 52588251 | *A1CF* | -1.19 | <0.00001 |
| cg02578087 | chr3 | 8671361 | *C3orf32* | -1.17 | <0.00001 |
| cg14612785 | chr3 | 71774842 | *EIF4E3* | -1.16 | <0.00001 |
| cg16628641 | chr16 | 4426265 | *VASN;CORO7* | -1.15 | <0.00001 |
| cg01861096 | chr22 | 19279212 | *CLTCL1* | -1.13 | <0.00001 |
| cg22855933 | chr2 | 157179512 | | -1.11 | <0.00001 |
| cg14435720 | chr9 | 139565087 | *EGFL7;MIR126* | -1.11 | <0.00001 |
| cg03885684 | chr2 | 120770471 | *EPB41L5* | 1.10 | <0.00001 |
| cg11849422 | chr9 | 112542571 | *PALM2;PALM2-AKAP2* | -1.10 | <0.00001 |
| cg16062483 | chr14 | 98444417 | *C14orf64* | -1.09 | <0.00001 |
| cg16150798 | chr11 | 61560092 | *MIR611;C11orf10;FEN1* | 1.08 | <0.00001 |
| cg08687137 | chr11 | 62389499 | *B3GAT3* | -1.08 | <0.00001 |
| cg18035158 | chr6 | 107810971 | *SOBP* | -1.07 | <0.00001 |
| cg21465162 | chr9 | 86323362 | *UBQLN1* | -1.06 | <0.00001 |
| cg13011999 | chr14 | 90167241 | | -1.06 | <0.00001 |
| cg17749261 | chr3 | 156392701 | *TIPARP-AS1;TIPARP* | -1.05 | <0.00001 |
| cg26506013 | chr5 | 133984553 | *SEC24A* | -1.04 | <0.00001 |

| | | | | | |
|---|---|---|---|---|---|
| cg10109500 | chr3 | 172165884 | *GHSR* | -1.04 | <0.00001 |
| cg13084560 | chr1 | 8418823 | *RERE* | -1.03 | <0.00001 |
| cg16786756 | chr4 | 187422114 | | -1.03 | <0.00001 |
| cg22984571 | chr3 | 186490656 | | -1.03 | <0.00001 |
| cg11581446 | chr9 | 99449206 | | 1.02 | <0.00001 |
| cg03567028 | chr13 | 44544293 | | -1.01 | <0.00001 |
| cg12524748 | chr4 | 1713366 | *SLBP* | -1.00 | <0.00001 |
| cg20124710 | chr15 | 55611105 | *PIGBOS1;PIGB;* | -1.00 | <0.00001 |
| cg03190876 | chr19 | 17566550 | *NXNL1* | -0.99 | <0.00001 |
| cg26870126 | chr10 | 124901477 | | 0.99 | <0.00001 |
| cg17491622 | chr1 | 1369616 | *VWA1* | -0.99 | <0.00001 |
| cg19375796 | chr3 | 52321643 | *GLYCTK* | -0.98 | <0.00001 |
| cg09428537 | chr16 | 90039076 | *AFG3L1;CENPBD1* | -0.98 | <0.00001 |
| cg25808577 | chr6 | 34433701 | *PACSIN1* | -0.98 | <0.00001 |
| cg14862827 | chr9 | 114936957 | *MIR3134;SUSD1* | -0.98 | <0.00001 |
| cg26674752 | chr10 | 43634212 | *CSGALNACT2* | -0.98 | <0.00001 |
| cg19308375 | chr2 | 228324937 | | -0.98 | <0.00001 |
| cg16222083 | chr1 | 212209131 | *DTL;INTS7* | -0.97 | <0.00001 |
| cg21223075 | chr3 | 5020327 | *BHLHE40* | 0.97 | <0.00001 |
| cg07360021 | chr6 | 151186904 | *MTHFD1L* | -0.96 | <0.00001 |
| cg00898988 | chr14 | 24020746 | *ZFHX2;THTPA* | -0.95 | <0.00001 |
| cg23616649 | chr12 | 49351112 | *ARF3* | -0.95 | <0.00001 |
| cg08186508 | chr14 | 68067006 | *PIGH* | -0.95 | <0.00001 |
| cg02294730 | chr15 | 41136779 | *SPINT1* | -0.95 | <0.00001 |
| cg05344293 | chr16 | 84627507 | *COTL1* | 0.94 | <0.00001 |
| cg19774733 | chr9 | 37422586 | *GRHPR* | -0.94 | <0.00001 |
| cg17592699 | chr4 | 3590676 | *LINC00955* | 0.94 | <0.00001 |
| cg01877318 | chr17 | 45266772 | *CDC27* | -0.94 | <0.00001 |
| cg00685135 | chr9 | 36136348 | *GLIPR2* | 0.91 | <0.00001 |
| cg01534423 | chr17 | 18965556 | | -0.90 | <0.00001 |
| cg21898346 | chr7 | 87505703 | *SLC25A40;DBF4* | -0.89 | <0.00001 |
| cg24237848 | chr1 | 32688013 | *TMEM234;EIF3I* | -0.89 | <0.00001 |

| | | | | | |
|---|---|---|---|---|---|
| cg14037413 | chr11 | 9482594 | *ZNF143* | -0.89 | <0.00001 |
| cg23850112 | chr3 | 40498671 | *RPL14* | -0.89 | <0.00001 |
| cg27139210 | chr14 | 37666963 | *MIPOL1* | -0.89 | <0.00001 |
| cg00088299 | chr4 | 2538934 | | 0.89 | <0.00001 |
| cg21855963 | chr1 | 78444801 | *FUBP1* | -0.88 | <0.00001 |
| cg07979357 | chr19 | 14142353 | *IL27RA* | -0.88 | <0.00001 |
| cg00542493 | chr10 | 12238021 | *NUDT5;CDC123* | -0.87 | <0.00001 |
| cg11998816 | chr12 | 56545925 | *MYL6B* | -0.87 | <0.00001 |
| cg02096758 | chr17 | 77394895 | *HRNBP3* | 0.87 | <0.00001 |
| cg17879791 | chr6 | 26217110 | *HIST1H2AE;HIST1H2BG* | -0.87 | <0.00001 |
| cg24017268 | chr1 | 6484816 | *ESPN* | -0.86 | <0.00001 |
| cg23633624 | chr8 | 95909079 | | -0.86 | <0.00001 |
| cg27041724 | chr9 | 20623705 | *MLLT3* | -0.85 | <0.00001 |
| cg27466766 | chr8 | 144679319 | *EEF1D;TIGD5* | 0.85 | <0.00001 |
| cg04802271 | chr22 | 39930989 | | 0.85 | <0.00001 |
| cg07731875 | chr20 | 43991546 | *SYS1;SYS1-DBNDD2* | -0.84 | <0.00001 |
| cg11208568 | chr2 | 122513714 | *TSN* | -0.84 | <0.00001 |
| cg01659459 | chr14 | 102227496 | *PPP2R5C* | 0.83 | <0.00001 |
| cg15578311 | chr3 | 58291756 | *RPP14* | -0.83 | <0.00001 |
| cg03834947 | chr15 | 65321822 | *MTFMT* | -0.83 | <0.00001 |
| cg16024729 | chr17 | 40224395 | | -0.82 | <0.00001 |
| cg25800379 | chr7 | 12250767 | *TMEM106B* | -0.82 | <0.00001 |
| cg22256960 | chr15 | 77711686 | | -0.81 | <0.00001 |
| cg22151281 | chr9 | 80850856 | *CEP78* | -0.81 | <0.00001 |
| cg05489143 | chr3 | 40498640 | *RPL14* | -0.81 | <0.00001 |
| cg10157938 | chr11 | 78757086 | *TENM4* | 0.81 | <0.00001 |
| cg00482802 | chr2 | 161126330 | *LOC100505984* | -0.80 | <0.00001 |
| cg15906479 | chr4 | 109089614 | *LEF1-AS1;LEF1* | -0.80 | <0.00001 |
| cg10532262 | chr3 | 172428426 | *NCEH1* | -0.80 | <0.00001 |
| cg03913456 | chr2 | 97000924 | *NCAPH* | -0.80 | <0.00001 |
| cg15801789 | chr9 | 128003861 | *HSPA5* | -0.80 | <0.00001 |
| cg06180910 | chr22 | 24382663 | *GSTT1* | -0.79 | <0.00001 |

| | | | | | |
|---|---|---|---|---|---|
| cg16595484 | chr3 | 122512170 | *HSPBAP1* | -0.79 | <0.00001 |
| cg14903832 | chr2 | 198318098 | *COQ10B* | -0.79 | <0.00001 |
| cg18493069 | chr6 | 108582623 | *SNX3* | -0.79 | <0.00001 |
| cg17036785 | chr12 | 110906113 | *FAM216A;GPN3* | -0.79 | <0.00001 |
| cg16542283 | chr4 | 76439453 | *THAP6;RCHY1* | -0.79 | <0.00001 |
| cg06561932 | chr8 | 23081981 | *LOC389641;TNFRSF10A* | -0.79 | <0.00001 |
| cg21995575 | chr2 | 47403831 | *CALM2* | -0.78 | <0.00001 |
| cg19781738 | chr1 | 52344674 | *NRD1* | -0.78 | <0.00001 |
| cg10722938 | chr19 | 14317606 | *LPHN1* | -0.77 | <0.00001 |
| cg08215925 | chr3 | 71633214 | *FOXP1* | -0.77 | <0.00001 |
| cg04883656 | chr6 | 71998597 | *OGFRL1* | -0.77 | <0.00001 |
| cg22533689 | chr1 | 16464479 | *EPHA2* | -0.77 | <0.00001 |
| cg19137818 | chr3 | 45837556 | *SLC6A20* | 0.77 | <0.00001 |
| cg20027289 | chr11 | 93861395 | *PANX1* | -0.77 | <0.00001 |
| cg13485746 | chr1 | 162467911 | *UHMK1* | -0.77 | <0.00001 |
| cg12538421 | chr8 | 133687986 | *LRRC6* | -0.77 | <0.00001 |
| cg11192793 | chr1 | 23886396 | *ID3* | -0.76 | <0.00001 |
| cg13831006 | chr7 | 128095730 | *HILPDA* | -0.76 | <0.00001 |
| cg10681804 | chr20 | 33433114 | *GGT7* | 0.76 | <0.00001 |
| cg00826921 | chr3 | 143692146 | *C3orf58* | -0.76 | <0.00001 |
| cg12791939 | chr19 | 6767670 | *SH2D3A* | -0.75 | <0.00001 |
| cg14322215 | chr6 | 28891358 | *TRIM27* | -0.75 | <0.00001 |
| cg14036868 | chr2 | 38604442 | *ATL2* | -0.75 | <0.00001 |
| cg23315685 | chr11 | 120081358 | *OAF* | -0.75 | <0.00001 |
| cg22655853 | chr3 | 112280943 | *ATG3;SLC35A5* | -0.75 | <0.00001 |
| cg01961410 | chr2 | 86790732 | *CHMP3;RNF103-CHMP3* | -0.75 | <0.00001 |
| cg13324603 | chr17 | 74733680 | *MFSD11;MIR636;SRSF2* | -0.74 | <0.00001 |
| cg04545708 | chr11 | 111750355 | *FDXACB1;C11orf1* | -0.74 | <0.00001 |
| cg23761970 | chr22 | 42765709 | *LINC01315* | -0.74 | <0.00001 |
| cg19642925 | chr10 | 103911966 | *NOLC1* | -0.73 | <0.00001 |
| cg17373554 | chr5 | 72594735 | | -0.73 | <0.00001 |
| cg04921814 | chr1 | 145575587 | *PIAS3* | -0.73 | <0.00001 |

| | | | | | |
|---|---|---|---|---|---|
| cg18106397 | chr12 | 29534230 | *ERGIC2* | -0.73 | <0.00001 |
| cg00461735 | chr1 | 234509013 | *C1orf31* | -0.73 | <0.00001 |
| cg12122453 | chr14 | 64971959 | *ZBTB25;ZBTB1* | -0.73 | <0.00001 |
| cg07119225 | chr9 | 99540647 | *ZNF510* | -0.73 | <0.00001 |
| cg25845597 | chr6 | 27841122 | *HIST1H4L;HIST1H3I* | -0.72 | <0.00001 |
| cg26133769 | chr4 | 30723855 | *PCDH7* | -0.72 | <0.00001 |
| cg19079372 | chr5 | 54604293 | *DHX29;SKIV2L2* | -0.71 | <0.00001 |
| cg06748143 | chr1 | 183387639 | *NMNAT2* | -0.71 | <0.00001 |
| cg16041806 | chr1 | 173991823 | | -0.71 | <0.00001 |
| cg17254222 | chr1 | 1981657 | *PRKCZ* | -0.71 | <0.00001 |
| cg16108521 | chr2 | 54013703 | *ERLEC1;ASB3;GPR75-ASB3* | -0.71 | <0.00001 |
| cg08965527 | chr16 | 84178213 | *HSDL1;LRRC50;HSDL1* | -0.71 | <0.00001 |
| cg23066318 | chr17 | 36860967 | *MLLT6* | -0.71 | <0.00001 |
| cg26954695 | chr19 | 59085561 | *MZF1;MGC2752;LOC100131691* | -0.70 | <0.00001 |
| cg16764909 | chr17 | 57233385 | *SKA2;PRR11* | -0.70 | <0.00001 |
| cg24891709 | chr7 | 138794209 | *ZC3HAV1* | -0.70 | <0.00001 |
| cg21807034 | chr18 | 61144150 | *SERPINB5* | 0.70 | <0.00001 |
| cg06638795 | chr2 | 42719933 | *KCNG3* | -0.69 | <0.00001 |

**Supplementary Table 3. Top 200 ranked probes meeting criteria for change (p<0.05 and >5%), and was unique to remitted ADHD.**

Chromosomal location (Chr), strand orientation, associated gene, and effect sizes (logFC) with unadjusted p-values provided.

| Probes | Chr | Position | *Gene* | logFC | p-value |
|---|---|---|---|---|---|
| cg04605287 | chr1 | 54953486 | | 0.996 | 0.0001 |
| cg25577322 | chr7 | 17338213 | *AHR* | 0.938 | 0.00001 |
| cg26199493 | chr1 | 226497589 | *LIN9* | -0.724 | 0.0001 |
| cg02482001 | chr17 | 6338476 | *AIPL1* | 0.338 | 0.02 |
| cg14824382 | chr8 | 144680115 | *EEF1D;TIGD5* | -0.331 | 0.02 |
| cg06921011 | chr10 | 102589467 | *PAX2* | 0.312 | 0.04 |
| cg08103988 | chr17 | 6558365 | | -0.302 | 0.04 |
| cg12077875 | chr20 | 47835964 | *DDX27* | 0.299 | 0.001 |
| cg09248826 | chr9 | 130860839 | *SLC25A25* | 0.297 | 0.003 |
| ch.14.64475309R | chr14 | 65405556 | | 0.297 | 0.001 |
| cg21944402 | chr4 | 77184893 | *FAM47E;FAM47E-STBD1* | -0.292 | <0.00001 |
| cg17154724 | chr1 | 171810322 | *DNM3* | 0.285 | 0.001 |
| cg13143743 | chr3 | 177570694 | | -0.278 | 0.0004 |
| cg05890377 | chr2 | 74357713 | | -0.277 | 0.005 |
| cg11641410 | chr8 | 54137552 | | -0.274 | 0.00001 |
| cg27377353 | chr8 | 81993869 | *PAG1* | 0.274 | 0.03 |
| cg22344617 | chr8 | 11152749 | *MTMR9* | -0.272 | 0.0003 |
| cg20257947 | chr4 | 167235409 | | -0.270 | 0.00001 |
| cg26665229 | chr8 | 75541549 | *MIR2052HG* | 0.270 | 0.00003 |
| cg11646117 | chr5 | 2100661 | | -0.269 | 0.0001 |
| cg14482748 | chr3 | 63075333 | | -0.269 | 0.00003 |
| cg24525457 | chr7 | 100091255 | *C7orf51* | 0.267 | 0.01 |
| cg10047753 | chr17 | 41438598 | | 0.266 | 0.005 |
| cg04608190 | chr2 | 240983080 | *PRR21* | -0.265 | 0.02 |
| cg17466535 | chr14 | 20929606 | *TMEM55B* | -0.264 | 0.004 |
| cg18489945 | chr15 | 60294398 | | 0.263 | 0.0003 |
| cg00975418 | chr9 | 129566028 | *ZBTB43* | -0.262 | 0.00002 |

| | | | | | |
|---|---|---|---|---|---|
| cg04774194 | chr6 | 127664674 | *ECHDC1* | -0.260 | <0.00001 |
| cg21813567 | chr7 | 79542782 | | -0.257 | 0.00004 |
| cg04733681 | chr6 | 160423822 | *IGF2R* | -0.257 | 0.002 |
| cg09692364 | chr13 | 76043069 | *TBC1D4* | -0.256 | 0.02 |
| cg13280056 | chr7 | 150103245 | *LOC728743* | -0.255 | 0.001 |
| cg06427702 | chr10 | 122228021 | *PPAPDC1A* | 0.254 | 0.01 |
| cg13409449 | chr10 | 8097354 | *GATA3* | 0.253 | 0.003 |
| cg27485596 | chr13 | 111829530 | *ARHGEF7* | 0.252 | 0.0001 |
| cg25730356 | chr1 | 154540159 | *CHRNB2* | 0.252 | 0.003 |
| cg03070989 | chr19 | 34311482 | | -0.250 | 0.01 |
| cg10439651 | chr1 | 161103008 | *DEDD* | 0.250 | 0.0003 |
| ch.18.700493F | chr18 | 35613416 | | 0.248 | 0.0004 |
| cg09521703 | chr19 | 55944864 | *SHISA7* | -0.247 | 0.02 |
| cg10557174 | chr13 | 29166467 | | 0.246 | 0.01 |
| cg20315445 | chr10 | 55393450 | | -0.244 | 0.0001 |
| cg17560015 | chr8 | 145013249 | *PLEC1* | 0.244 | 0.0002 |
| cg11468993 | chr6 | 31082939 | *CDSN;PSORS1C1* | 0.243 | 0.01 |
| cg08078339 | chr2 | 43848123 | | -0.243 | 0.00003 |
| cg25956844 | chr3 | 179370364 | *USP13* | 0.243 | 0.00002 |
| cg02045285 | chr17 | 36861674 | *MLLT6* | -0.243 | 0.00003 |
| cg04688351 | chr2 | 223154140 | *PAX3* | 0.243 | 0.001 |
| cg01778565 | chr8 | 115855565 | | -0.242 | 0.0001 |
| ch.13.639148F | chr13 | 47925785 | | 0.241 | 0.00003 |
| cg27153400 | chr19 | 55973320 | *ISOC2* | 0.241 | 0.01 |
| cg05401447 | chr7 | 100728731 | *TRIM56* | 0.241 | 0.002 |
| cg05962718 | chr1 | 93545112 | *MTF2* | 0.241 | 0.001 |
| cg18678645 | chr5 | 135416331 | *MIR886* | 0.241 | 0.02 |
| cg08599031 | chr8 | 86020704 | *LRRCC1* | -0.240 | 0.0001 |
| cg03674660 | chr6 | 168844698 | *SMOC2* | -0.240 | 0.00003 |
| cg08787030 | chr5 | 97803704 | | -0.240 | 0.001 |
| cg18087943 | chr11 | 2160540 | *INS-IGF2;IGF2AS;IGF2* | -0.240 | 0.0004 |
| cg00261149 | chr5 | 180633181 | *TRIM7* | 0.240 | 0.001 |

| | | | | | |
|---|---|---|---|---|---|
| cg01019631 | chr10 | 112436302 | *RBM20* | 0.239 | 0.02 |
| cg22728128 | chr1 | 112299491 | *DDX20;C1orf183* | -0.239 | 0.0003 |
| cg26031255 | chr6 | 150463965 | *PPP1R14C* | 0.239 | 0.001 |
| cg15338778 | chr6 | 116574713 | *TSPYL4* | 0.239 | 0.0001 |
| cg23276602 | chr2 | 55617438 | *CCDC88A* | -0.238 | 0.004 |
| cg19103429 | chr7 | 92460784 | *CDK6* | 0.238 | 0.02 |
| cg21359576 | chr12 | 109124717 | *CORO1C* | 0.238 | 0.00001 |
| cg23616258 | chr7 | 95106564 | | 0.238 | 0.0002 |
| cg08447739 | chr10 | 124220359 | *HTRA1* | -0.237 | 0.01 |
| cg15025532 | chr4 | 39460522 | *RPL9;LIAS* | 0.237 | 0.00001 |
| cg22974158 | chr10 | 25305096 | *THNSL1;ENKUR* | 0.237 | 0.001 |
| cg00909993 | chr2 | 120516537 | *PTPN4* | -0.236 | 0.00004 |
| cg14610566 | chr2 | 101086506 | *NMS* | -0.236 | 0.001 |
| cg11956562 | chr9 | 33983377 | *UBAP2* | -0.236 | 0.0004 |
| cg17538538 | chr6 | 106825021 | | 0.236 | 0.0001 |
| cg05041045 | chr14 | 31091339 | *SCFD1* | 0.236 | 0.0005 |
| cg14355192 | chr9 | 36258114 | *GNE* | 0.235 | 0.0002 |
| cg26942031 | chr6 | 163912396 | *QKI* | -0.235 | 0.02 |
| cg18095725 | chr6 | 26159136 | *HIST1H2BD* | 0.234 | 0.0001 |
| cg21007691 | chr14 | 102695264 | *RAGE* | -0.233 | 0.0003 |
| ch.8.1995451R | chr8 | 98920744 | *MATN2* | 0.233 | 0.002 |
| cg19207921 | chr4 | 148451778 | *EDNRA* | -0.232 | 0.0004 |
| cg10211209 | chr13 | 43410862 | | -0.231 | 0.00001 |
| cg01204603 | chr16 | 80619852 | | -0.231 | 0.0002 |
| cg21051694 | chr9 | 114724064 | | 0.231 | 0.00007 |
| cg11126410 | chr3 | 128841191 | *RAB43;ISY1-RAB43* | -0.231 | 0.002 |
| cg03637878 | chr11 | 133938788 | *JAM3* | 0.230 | 0.001 |
| cg19736503 | chr6 | 84563604 | *RIPPLY2* | 0.230 | 0.001 |
| cg08637269 | chr11 | 122524689 | | -0.230 | 0.05 |
| cg25703407 | chr12 | 69725223 | | 0.230 | 0.001 |
| cg09430341 | chr1 | 46090209 | *CCDC17* | -0.230 | 0.02 |
| cg11911679 | chr2 | 182520919 | *CERKL* | -0.229 | 0.00002 |

| | | | | | |
|---|---|---|---|---|---|
| cg06673020 | chr15 | 95974945 | *LINC00924* | -0.229 | 0.0001 |
| cg02134923 | chr2 | 42193753 | | -0.229 | 0.03 |
| cg02571311 | chr15 | 99813283 | *LRRC28* | 0.229 | 0.0004 |
| cg17985493 | chr1 | 25102228 | *CLIC4* | -0.229 | 0.00003 |
| cg11947509 | chr2 | 202898349 | *FZD7* | -0.229 | 0.0002 |
| cg22982767 | chr22 | 46454012 | *LOC150381* | 0.229 | 0.001 |
| cg18934287 | chr13 | 46967579 | | -0.228 | 0.00002 |
| cg11304315 | chr4 | 109541595 | *RPL34;LOC285456* | 0.228 | 0.0004 |
| cg02023043 | chr7 | 45129337 | *NACAD* | 0.228 | 0.001 |
| cg08845486 | chr16 | 67217724 | *KIAA0895L* | 0.227 | 0.0001 |
| cg13749379 | chr17 | 41833161 | *SOST* | -0.227 | 0.01 |
| cg16780862 | chr19 | 2346131 | *SPPL2B* | -0.226 | 0.0003 |
| cg26873042 | chr5 | 67363338 | | 0.226 | 0.01 |
| cg00457246 | chr1 | 243904416 | *AKT3* | -0.226 | 0.0001 |
| cg22078934 | chr2 | 28905057 | | -0.226 | <0.00001 |
| cg19942383 | chr3 | 42979226 | | -0.226 | 0.01 |
| cg26128378 | chr8 | 1585146 | *DLGAP2* | -0.225 | 0.00001 |
| cg11641097 | chr4 | 42054953 | *SLC30A9* | -0.225 | 0.004 |
| cg06743283 | chr19 | 58382549 | *ZNF814* | -0.225 | 0.0002 |
| cg04571941 | chr7 | 5463284 | *TNRC18* | -0.225 | <0.00001 |
| ch.2.4469412F | chr2 | 224066581 | | 0.224 | 0.002 |
| cg23141147 | chr19 | 2962227 | | 0.224 | 0.0003 |
| cg09215316 | chr12 | 7033628 | *ATN1* | 0.224 | 0.0004 |
| cg04959674 | chr10 | 99257608 | *UBTD1;MMS19* | 0.224 | 0.002 |
| cg07528209 | chr2 | 121549725 | | -0.223 | 0.02 |
| cg16280579 | chr2 | 96965285 | *SNRNP200* | -0.223 | 0.0003 |
| cg05229528 | chr11 | 30409857 | *MPPED2* | -0.223 | 0.001 |
| cg09982326 | chr11 | 94568871 | *AMOTL1* | -0.223 | 0.003 |
| cg15487187 | chr22 | 22555410 | | 0.223 | 0.03 |
| cg12011711 | chr17 | 65496598 | *PITPNC1* | -0.222 | 0.003 |
| cg21514373 | chr1 | 25524532 | | 0.222 | 0.001 |
| cg26601775 | chr6 | 42185256 | *MRPS10* | 0.222 | 0.0005 |

| | | | | | |
|---|---|---|---|---|---|
| cg05948408 | chr5 | 88178572 | *MEF2C* | 0.222 | 0.002 |
| cg09720597 | chr2 | 111907136 | *BCL2L11* | -0.221 | 0.0001 |
| cg24873093 | chr1 | 45991060 | | 0.221 | 0.01 |
| cg16434720 | chr7 | 130793000 | *LINC-PINT* | 0.221 | 0.0002 |
| cg08045906 | chr6 | 32120625 | *PPT2;PRRT1* | -0.221 | 0.04 |
| cg09630103 | chr14 | 75868204 | | -0.221 | 0.02 |
| cg06023901 | chr6 | 57037482 | *BAG2* | 0.221 | 0.00004 |
| cg17509172 | chr3 | 152552215 | *P2RY1* | 0.220 | 0.004 |
| cg04170952 | chr17 | 33957779 | *AP2B1* | -0.220 | 0.01 |
| cg05810439 | chr4 | 15608515 | *FBXL5* | -0.220 | 0.0002 |
| cg22888160 | chr1 | 152177273 | | 0.220 | 0.02 |
| cg08356083 | chr16 | 27280211 | *NSMCE1* | 0.219 | 0.0002 |
| cg20796179 | chr9 | 123837475 | | -0.219 | 0.01 |
| cg10305928 | chr10 | 62426219 | *ANK3* | 0.219 | 0.03 |
| cg10961853 | chr10 | 77172915 | | -0.218 | 0.0003 |
| cg14687930 | chr13 | 113125773 | | -0.218 | 0.0004 |
| cg09665331 | chr1 | 195867753 | | 0.218 | 0.01 |
| cg05863502 | chr9 | 140771990 | *CACNA1B* | -0.217 | 0.0001 |
| cg24663419 | chr2 | 58862141 | | -0.217 | 0.01 |
| cg23635910 | chr1 | 912055 | *C1orf170* | -0.217 | 0.01 |
| cg26402284 | chr14 | 105500294 | | -0.217 | 0.0001 |
| cg10905593 | chr10 | 102027479 | *CWF19L1* | 0.217 | 0.002 |
| cg00924460 | chr11 | 32175112 | | 0.216 | 0.03 |
| cg02313132 | chr16 | 74971138 | *WDR59* | 0.216 | 0.04 |
| cg20473723 | chr20 | 6104884 | *FERMT1* | -0.216 | 0.002 |
| cg23581650 | chr10 | 120551894 | | -0.216 | 0.01 |
| cg03787899 | chr5 | 661046 | *TPPP* | -0.215 | 0.0008 |
| cg21571793 | chr14 | 89910404 | *FOXN3* | -0.215 | 0.001 |
| cg21187830 | chr3 | 44552297 | *ZNF852* | 0.215 | 0.002 |
| cg10805254 | chr3 | 72433837 | *RYBP* | 0.215 | 0.03 |
| cg11034073 | chr4 | 88719777 | *IBSP* | -0.215 | 0.00001 |
| cg26433494 | chr7 | 132068972 | *PLXNA4* | -0.215 | 0.01 |

| | | | | | |
|---|---|---|---|---|---|
| cg13564087 | chr19 | 56825799 | | 0.215 | 0.00005 |
| cg11789421 | chr10 | 103543213 | *NPM3* | 0.215 | 0.001 |
| cg23396843 | chr7 | 97821646 | *LMTK2* | -0.214 | 0.0003 |
| cg11495303 | chr6 | 26024165 | | -0.214 | 0.00003 |
| ch.8.614314F | chr8 | 25064094 | *DOCK5* | 0.214 | 0.0003 |
| cg17966245 | chr8 | 95523925 | *KIAA1429* | -0.214 | 0.002 |
| cg23321751 | chr18 | 34429726 | *KIAA1328* | -0.213 | 0.01 |
| cg16600869 | chr1 | 40349633 | *TRIT1* | -0.213 | 0.0001 |
| cg14963901 | chr18 | 30471820 | | -0.213 | 0.01 |
| cg06123563 | chr10 | 4981518 | | -0.213 | 0.05 |
| cg23353374 | chr4 | 15693171 | | 0.213 | 0.01 |
| cg20460697 | chr18 | 56338298 | *MALT1* | 0.213 | 0.04 |
| cg26516362 | chr5 | 178986906 | *RUFY1* | -0.213 | 0.04 |
| cg16293575 | chr11 | 6948211 | *ZNF215* | 0.213 | 0.001 |
| cg26065130 | chr2 | 100023445 | *REV1* | -0.213 | 0.001 |
| cg23753855 | chr5 | 88179010 | *MEF2C;MEF2C-AS1* | 0.213 | 0.01 |
| cg08528486 | chr13 | 113648767 | *MCF2L* | -0.212 | 0.01 |
| cg07917289 | chr19 | 45147056 | *PVR* | 0.212 | 0.0001 |
| cg21433912 | chr7 | 84815088 | | 0.212 | 0.0005 |
| cg26367649 | chr6 | 91297042 | *MAP3K7* | 0.212 | 0.002 |
| cg00046410 | chr1 | 35581062 | *ZMYM1* | -0.212 | 0.0001 |
| cg17683773 | chr15 | 78634553 | *CRABP1* | -0.212 | 0.0003 |
| cg19812018 | chr17 | 55824035 | *CCDC182* | -0.212 | 0.001 |
| cg10566589 | chr6 | 111927040 | *TRAF3IP2* | 0.212 | 0.001 |
| cg11134825 | chr1 | 90185796 | | 0.212 | 0.02 |
| cg05179986 | chr6 | 151547145 | | -0.212 | 0.002 |
| cg03850986 | chr10 | 116408382 | *ABLIM1* | 0.212 | 0.003 |
| cg11270956 | chr1 | 184048238 | | -0.212 | <0.00001 |
| cg06614611 | chr12 | 115221608 | | -0.211 | 0.0002 |
| cg04529582 | chr20 | 30697510 | *TM9SF4* | 0.211 | 0.001 |
| cg06965316 | chr10 | 72163829 | *EIF4EBP2* | -0.211 | 0.002 |
| cg20927395 | chr3 | 115120356 | | 0.211 | 0.0003 |

| | | | | | |
|---|---|---|---|---|---|
| cg02781074 | chr13 | 101241206 | *GGACT* | -0.211 | 0.01 |
| cg12716232 | chr8 | 43001931 | *HGSNAT* | -0.210 | 0.00003 |
| cg13716787 | chr2 | 70351227 | *LOC100133985* | 0.210 | 0.0002 |
| cg09732156 | chr14 | 54349470 | | -0.210 | 0.0002 |
| cg16713996 | chr5 | 112202687 | *SRP19* | -0.210 | 0.02 |
| cg12037340 | chr10 | 94812063 | *EXOC6* | 0.210 | 0.001 |
| cg11459237 | chr6 | 111029441 | *CDK19* | -0.210 | 0.0001 |
| cg01317772 | chr16 | 55405200 | | 0.210 | 0.01 |
| cg11527760 | chr4 | 107173212 | *TBCK* | -0.210 | 0.0002 |
| cg05086789 | chr10 | 29701805 | *LOC387647* | -0.210 | 0.0002 |
| cg22998101 | chr11 | 55763230 | *OR5F1* | -0.209 | 0.003 |
| cg07915976 | chr12 | 54414427 | *HOXC4;HOXC5;HOXC6* | -0.209 | 0.001 |
| cg21020089 | chr6 | 41863578 | *USP49* | 0.209 | 0.0002 |

**Supplementary Table 4. Top 200 differentially methylated probes between ADHD and controls. Top ranked CpG probes showing significant main effects (n=100) or diagnosis x time interactions (n=100) in comparison between pooled ADHD and control groups.** Chromosomal location (Chr), strand orientation, associated gene, and effect sizes (logFC) with unadjusted p-values provided.

| Probes | Chr | Position | *Gene* | logFC | p-value |
|---|---|---|---|---|---|
| *Main effect* | | | | | |
| cg17217665 | chr6 | 44186914 | *SLC29A1* | 0.36 | <0.00001 |
| cg07817266 | chr17 | 43765890 | *MGC57346-CRHR1* | 0.37 | <0.00001 |
| cg18401534 | chr7 | 5710286 | *RNF216-IT1;RNF216;RNF216* | -0.32 | 0.00001 |
| cg17628588 | chr16 | 3580633 | *CLUAP1* | -0.23 | 0.00001 |
| cg12881363 | chr22 | 50528298 | *MOV10L1* | 0.82 | 0.00001 |
| cg05641033 | chr12 | 1639534 | | 0.34 | 0.00001 |
| cg00684531 | chr4 | 1994019 | *NELFA* | -0.41 | 0.00001 |
| cg12577151 | chr3 | 128615097 | *ACAD9* | -0.29 | 0.00001 |
| cg01206832 | chr1 | 178622333 | | -0.24 | 0.00002 |
| cg02062409 | chr12 | 98850881 | | 0.38 | 0.00002 |
| cg01850934 | chr14 | 68141664 | *VTI1B* | -0.33 | 0.00002 |
| cg13353337 | chr22 | 21336550 | *LZTR1* | -0.83 | 0.00003 |
| cg04616793 | chr16 | 8738608 | *C16orf68* | -0.23 | 0.00003 |
| cg01525538 | chr1 | 228785987 | *DUSP5P* | 0.32 | 0.00003 |
| cg09335159 | chr11 | 129147936 | | -0.26 | 0.00003 |
| cg10636959 | chr6 | 30162942 | *TRIM26* | 0.25 | 0.00003 |
| cg04357253 | chr12 | 110662671 | | -0.32 | 0.00004 |
| cg01722420 | chr15 | 32993763 | | -0.28 | 0.00004 |
| cg05164926 | chr17 | 7255624 | *KCTD11* | 0.44 | 0.00004 |
| cg08584037 | chr9 | 4984071 | *JAK2* | 0.27 | 0.00004 |
| cg12878847 | chr10 | 112515808 | *RBM20* | -0.32 | 0.00005 |
| cg15550572 | chr1 | 2064441 | *PRKCZ* | -0.48 | 0.00005 |
| cg12708331 | chr1 | 3440961 | *MEGF6* | -0.26 | 0.00005 |
| cg04438997 | chr17 | 70115868 | *SOX9* | 0.22 | 0.0001 |

| | | | | | |
|---|---|---|---|---|---|
| cg23364517 | chr15 | 86623057 | | -0.31 | 0.0001 |
| cg26939377 | chr19 | 41869719 | *B9D2*;*TMEM91* | -0.52 | 0.0001 |
| cg22760019 | chr3 | 136559194 | *SLC35G2* | -0.25 | 0.0001 |
| cg03080336 | chr4 | 186033378 | | -0.22 | 0.0001 |
| cg11124426 | chr2 | 131149108 | | -0.41 | 0.0001 |
| cg25602485 | chr1 | 234611287 | *TARBP1* | 0.28 | 0.0001 |
| cg22821026 | chr1 | 155108104 | *RAG1AP1* | 0.27 | 0.0001 |
| cg05677467 | chr15 | 85197654 | *WDR73* | -0.32 | 0.0001 |
| cg01756841 | chr1 | 176176579 | *RFWD2* | -0.21 | 0.0001 |
| cg01942468 | chr16 | 58233183 | *CSNK2A2* | -0.30 | 0.0001 |
| cg02853616 | chr4 | 94755787 | | 0.32 | 0.0001 |
| cg24273599 | chr10 | 79542494 | | -0.27 | 0.0001 |
| cg00934611 | chr2 | 175351869 | *GPR155* | -0.44 | 0.0001 |
| cg02692522 | chr18 | 61525844 | | -0.51 | 0.0001 |
| cg10808399 | chr17 | 7297336 | *PLSCR3* | 0.22 | 0.0001 |
| cg19646227 | chr1 | 238081674 | *LOC100130331* | 0.25 | 0.0001 |
| cg17446824 | chr7 | 150929507 | *CHPF2* | 0.34 | 0.0001 |
| cg21989213 | chr17 | 41984470 | *MPP2* | -0.44 | 0.0001 |
| cg15613047 | chr7 | 62557964 | | 0.21 | 0.0001 |
| cg09617296 | chr13 | 100153694 | *TM9SF2* | -0.36 | 0.0001 |
| cg25564901 | chr17 | 17140463 | *FLCN* | -0.28 | 0.0001 |
| cg03996398 | chr11 | 128813442 | *TP53AIP1* | -0.19 | 0.0001 |
| cg12224508 | chr2 | 28589782 | | 0.23 | 0.0001 |
| cg12609785 | chr17 | 43660871 | | 0.48 | 0.0001 |
| cg10831479 | chr17 | 61501560 | *TANC2* | -0.23 | 0.0001 |
| cg14693194 | chr18 | 6415252 | *L3MBTL4* | -0.18 | 0.0001 |
| cg14884987 | chr10 | 5496232 | *NET1* | -0.20 | 0.0001 |
| cg16597406 | chr17 | 8130173 | *C17orf68* | 0.36 | 0.0001 |
| cg01722566 | chr16 | 84975714 | | -0.25 | 0.0001 |
| cg22497095 | chr6 | 28601375 | | -0.39 | 0.0001 |
| cg06306684 | chr11 | 911512 | *CHID1* | -0.44 | 0.0001 |
| cg21210555 | chr14 | 69287543 | | 0.47 | 0.0001 |

| | | | | | |
|---|---|---|---|---|---|
| cg14819132 | chr17 | 17495032 | *PEMT* | -0.38 | 0.0001 |
| cg23238440 | chr10 | 124222816 | *HTRA1* | -0.19 | 0.0001 |
| cg12410530 | chr22 | 32001086 | *SFI1* | -0.55 | 0.0001 |
| cg15148757 | chr4 | 87857305 | *AFF1* | -0.31 | 0.0001 |
| cg02271621 | chr11 | 1592675 | *HCCA2*;*LOC338651*;*DUSP8* | 0.38 | 0.0001 |
| cg00734838 | chr10 | 14614164 | *FAM107B* | 0.38 | 0.0001 |
| cg12259302 | chr15 | 45958400 | *SQRDL* | -0.21 | 0.0001 |
| cg09590401 | chr12 | 132106088 | | -0.18 | 0.0001 |
| cg15649852 | chr7 | 65879115 | | -0.45 | 0.0001 |
| cg18145578 | chr2 | 64470508 | *LOC100507006* | -0.20 | 0.0001 |
| cg00678472 | chr3 | 44941712 | *TGM4* | -0.46 | 0.0002 |
| cg22914188 | chr12 | 110841667 | *ANAPC7* | -0.48 | 0.0002 |
| cg02597966 | chr5 | 125345473 | | 0.29 | 0.0002 |
| cg10533057 | chr4 | 152494843 | *FAM160A1* | -0.23 | 0.0002 |
| cg16421413 | chr2 | 47195558 | *TTC7A* | -0.23 | 0.0002 |
| cg08564601 | chr11 | 60048221 | *MS4A4A* | 0.23 | 0.0002 |
| cg08983990 | chr1 | 111682957 | *DRAM2;CEPT1* | -0.35 | 0.0002 |
| cg18728732 | chr6 | 36349532 | *ETV7* | 0.22 | 0.0002 |
| cg01864498 | chr10 | 81892291 | *PLAC9* | 0.43 | 0.0002 |
| cg13250661 | chr3 | 151960339 | | -0.30 | 0.0002 |
| cg18508935 | chr2 | 20307733 | | -0.24 | 0.0002 |
| cg13297150 | chr2 | 74710531 | *TTC31*;*CCDC142* | -0.34 | 0.0002 |
| cg22338845 | chr11 | 123429542 | *GRAMD1B* | -0.28 | 0.0002 |
| cg18444875 | chr8 | 107460168 | *OXR1* | 0.33 | 0.0002 |
| cg18059362 | chr5 | 149880708 | *NDST1* | -0.32 | 0.0002 |
| cg15824752 | chr1 | 171291393 | *FMO4* | -0.24 | 0.0002 |
| cg05192589 | chr1 | 44440480 | *ATP6V0B* | 0.31 | 0.0002 |
| cg08650016 | chr11 | 62359048 | *TUT1* | -0.41 | 0.0002 |
| cg24273709 | chr12 | 56308490 | *WIBG* | -0.26 | 0.0002 |
| cg07718702 | chr2 | 98800985 | *VWA3B* | -0.18 | 0.0002 |
| cg12079740 | chr1 | 240473071 | *FMN2* | -0.18 | 0.0002 |
| cg02911248 | chr2 | 62678372 | | -0.25 | 0.0002 |

| | | | | | |
|---|---|---|---|---|---|
| cg02530971 | chr2 | 47169235 | *MCFD2;TTC7A* | -0.30 | 0.0002 |
| cg14834568 | chr10 | 14614421 | *FAM107B* | 0.43 | 0.0002 |
| cg27625802 | chr4 | 4396613 | *NSG1* | -0.29 | 0.0002 |
| cg26344227 | chr7 | 53287126 | | 0.41 | 0.0002 |
| cg21156793 | chr2 | 71301576 | *NAGK* | -0.26 | 0.0002 |
| cg02426324 | chr7 | 40680454 | *C7orf10* | 0.26 | 0.0002 |
| cg09737668 | chr1 | 159923525 | *SLAMF9* | -0.19 | 0.0002 |
| cg12623640 | chr20 | 30552396 | | -0.14 | 0.0002 |
| cg19690214 | chr6 | 154678326 | *IPCEF1* | -0.24 | 0.0002 |
| cg05923197 | chr14 | 54418804 | *BMP4* | 0.19 | 0.0002 |
| cg03759229 | chr6 | 28601417 | | -0.32 | 0.0002 |
| cg26357405 | chr3 | 119296959 | *ADPRH* | 0.31 | 0.0003 |
| *Interaction effect* | | | | | |
| cg05600164 | chr8 | 38259904 | *LETM2* | 0.21 | <0.00001 |
| cg26901352 | chr16 | 57447502 | *CCL17* | -0.10 | <0.00001 |
| cg10636959 | chr6 | 30162942 | *TRIM26* | -0.13 | <0.00001 |
| cg20924302 | chr14 | 91156070 | *TTC7B* | 0.14 | 0.00001 |
| cg13250661 | chr3 | 151960339 | | 0.17 | 0.00001 |
| cg18225895 | chr6 | 52149972 | *MCM3* | 0.18 | 0.00001 |
| cg13318817 | chr2 | 121955116 | | 0.19 | 0.00001 |
| cg14693194 | chr18 | 6415252 | *L3MBTL4* | 0.10 | 0.00001 |
| cg11258630 | chr9 | 98857209 | *LOC158435* | -0.12 | 0.00001 |
| cg09351156 | chr6 | 18436845 | *RNF144B* | 0.16 | 0.00001 |
| cg25044635 | chr2 | 25565267 | *DNMT3A* | -0.20 | 0.00001 |
| cg18506547 | chr14 | 67956205 | *TMEM229B* | -0.12 | 0.00002 |
| cg02780269 | chr17 | 80292252 | *SECTM1* | -0.20 | 0.00002 |
| cg20955020 | chr21 | 28215410 | *ADAMTS1* | 0.13 | 0.00003 |
| cg25836301 | chr14 | 101292306 | *MEG3* | -0.10 | 0.00003 |
| cg03243362 | chr1 | 32741543 | *LCK* | 0.14 | 0.00003 |
| cg18035914 | chr16 | 81070732 | *ATMIN* | 0.17 | 0.00003 |
| cg02970775 | chr9 | 140483477 | *ZMYND19* | 0.15 | 0.00003 |
| cg09153256 | chr15 | 78977915 | | 0.19 | 0.00004 |

| | | | | | |
|---|---|---|---|---|---|
| cg21498648 | chr1 | 28453832 | | 0.13 | 0.00005 |
| cg14839558 | chr1 | 1874488 | | -0.13 | 0.00005 |
| cg05590862 | chr4 | 184093509 | *WWC2* | -0.11 | 0.00005 |
| cg25444339 | chr7 | 75194698 | *HIP1* | 0.18 | 0.0001 |
| cg09900463 | chr16 | 3179627 | | 0.18 | 0.0001 |
| cg06374962 | chr4 | 111531926 | | -0.11 | 0.0001 |
| cg20077208 | chr11 | 65349222 | *EHBP1L1* | -0.09 | 0.0001 |
| cg18755829 | chr1 | 38512230 | *POU3F1* | -0.14 | 0.0001 |
| cg17240415 | chr12 | 91755459 | | 0.14 | 0.0001 |
| cg20138458 | chr18 | 74110979 | *ZNF516* | 0.17 | 0.0001 |
| cg11199798 | chr11 | 10525508 | *AMPD3* | 0.16 | 0.0001 |
| cg00680728 | chr21 | 38417649 | | -0.10 | 0.0001 |
| cg00705004 | chr8 | 6827814 | *DEFA10P* | -0.13 | 0.0001 |
| cg13297150 | chr2 | 74710531 | *TTC31;CCDC142* | 0.17 | 0.0001 |
| cg24926042 | chr12 | 52799999 | *KRT82* | -0.10 | 0.0001 |
| cg16261511 | chr1 | 42471958 | *HIVEP3* | -0.15 | 0.0001 |
| cg08224563 | chr16 | 20916305 | *LYRM1* | 0.17 | 0.0001 |
| cg12118082 | chr3 | 133748785 | *SLCO2A1* | -0.16 | 0.0001 |
| cg08091310 | chr14 | 100989105 | *WDR25* | 0.08 | 0.0001 |
| cg18064927 | chr4 | 111531975 | | -0.14 | 0.0001 |
| cg20263853 | chr1 | 8824177 | *RERE* | -0.10 | 0.0001 |
| cg09838956 | chr15 | 25325713 | *SNORD116-15* | -0.10 | 0.0001 |
| cg03083227 | chr16 | 16235187 | *ABCC1* | -0.10 | 0.0001 |
| cg22389654 | chr1 | 120273819 | *PHGDH* | 0.10 | 0.0001 |
| cg08764599 | chr12 | 12620523 | | -0.12 | 0.0001 |
| cg05585630 | chr7 | 157510462 | *PTPRN2* | -0.15 | 0.0001 |
| cg13764927 | chr6 | 168553516 | | 0.10 | 0.0001 |
| cg24343720 | chr21 | 31539057 | *CLDN17* | 0.13 | 0.0001 |
| cg18855165 | chr9 | 93959685 | | 0.19 | 0.0001 |
| cg25842050 | chr5 | 126181581 | | 0.17 | 0.0001 |
| cg18788040 | chr3 | 128952552 | | 0.09 | 0.0001 |
| cg22906400 | chr1 | 11550802 | *PTCHD2* | -0.10 | 0.0001 |

| | | | | | |
|---|---|---|---|---|---|
| cg16899444 | chr3 | 151119739 | *MED12L* | -0.11 | 0.0001 |
| cg10343639 | chr1 | 153751665 | *SLC27A3* | 0.08 | 0.0001 |
| cg03386156 | chr18 | 66381799 | *CCDC102B*;*TMX3* | 0.12 | 0.0001 |
| cg13612623 | chr17 | 28679114 | | 0.14 | 0.0001 |
| cg16090273 | chr5 | 149737303 | *TCOF1* | -0.12 | 0.0001 |
| cg03071553 | chr19 | 5867086 | *FUT5* | 0.11 | 0.0001 |
| cg25480437 | chr1 | 159722028 | | 0.20 | 0.0001 |
| cg08174386 | chr2 | 1550610 | | -0.10 | 0.0001 |
| cg05853522 | chr11 | 65278442 | | 0.14 | 0.0001 |
| cg17996835 | chr13 | 111145409 | *COL4A2* | -0.10 | 0.0001 |
| cg04867484 | chr4 | 149297469 | *NR3C2* | 0.16 | 0.0001 |
| cg05903707 | chr2 | 158843896 | | 0.08 | 0.0001 |
| cg26656452 | chr10 | 115313165 | *HABP2* | 0.10 | 0.0002 |
| cg00125406 | chr4 | 41937906 | *TMEM33* | 0.12 | 0.0002 |
| cg04610108 | chr7 | 45140740 | *TBRG4* | -0.10 | 0.0002 |
| cg25845747 | chr6 | 57037243 | *BAG2* | -0.16 | 0.0002 |
| cg00314230 | chr21 | 45759016 | *C21orf2* | -0.13 | 0.0002 |
| cg20536716 | chr7 | 119913368 | *KCND2* | -0.18 | 0.0002 |
| cg18401534 | chr7 | 5710286 | *RNF216-IT1*;*RNF216* | 0.13 | 0.0002 |
| cg16506703 | chr16 | 75529356 | *CHST6* | -0.14 | 0.0002 |
| cg25121841 | chr15 | 93665609 | | -0.09 | 0.0002 |
| cg15112081 | chr3 | 183973581 | *ECE2* | 0.10 | 0.0002 |
| cg13151645 | chr15 | 102196103 | *TARSL2* | -0.10 | 0.0002 |
| cg25048987 | chr12 | 8645990 | | 0.14 | 0.0002 |
| cg01755336 | chr14 | 102653321 | *WDR20* | 0.09 | 0.0002 |
| cg01919429 | chr13 | 100154415 | *TM9SF2* | 0.11 | 0.0002 |
| cg13911707 | chr1 | 247496404 | *ZNF496* | -0.11 | 0.0002 |
| cg16223785 | chr11 | 8693620 | | 0.09 | 0.0002 |
| cg03576976 | chr7 | 155352634 | | -0.09 | 0.0002 |
| cg21954824 | chr1 | 151451556 | | 0.08 | 0.0002 |
| cg00011461 | chr12 | 57432410 | *MYO1A* | -0.11 | 0.0002 |
| cg11974931 | chr4 | 40632880 | *RBM47* | 0.14 | 0.0002 |

| | | | | | |
|---|---|---|---|---|---|
| cg02090806 | chr6 | 33239496 | *VPS52*;*RPS18* | 0.20 | 0.0002 |
| cg09141216 | chr15 | 48483907 | *CTXN2* | 0.17 | 0.0002 |
| cg03239925 | chr18 | 77230795 | *NFATC1* | -0.15 | 0.0002 |
| cg07293314 | chr12 | 109012166 | | 0.16 | 0.0002 |
| cg20693099 | chr11 | 76912768 | *MYO7A* | -0.11 | 0.0002 |
| cg22891632 | chr20 | 57184191 | *APCDD1L-AS1* | -0.12 | 0.0002 |
| cg15260925 | chr3 | 71436970 | *FOXP1* | 0.15 | 0.0002 |
| cg04787928 | chr3 | 119318672 | *PLA1A* | -0.14 | 0.0002 |
| cg16720242 | chr1 | 46196138 | *IPP* | 0.16 | 0.0002 |
| cg02919982 | chr1 | 153005121 | *SPRR1B* | -0.07 | 0.0002 |
| cg03482769 | chr19 | 10445593 | *RAVER1*;*ICAM3* | -0.17 | 0.0002 |
| cg22370231 | chr14 | 88462612 | | 0.13 | 0.0002 |
| cg20707195 | chr2 | 25596386 | | -0.13 | 0.0002 |
| cg23404330 | chr17 | 80251639 | | -0.17 | 0.0002 |
| cg05412784 | chr8 | 68379155 | *CPA6* | -0.14 | 0.0002 |
| cg23154281 | chr8 | 102517078 | *GRHL2* | 0.22 | 0.0002 |
| cg25714069 | chr22 | 20791214 | *SCARF2* | -0.15 | 0.0002 |

**Supplementary Table 5. Top 20 ranked GO and KEGG pathway terms linked to 200 differentially methylation probes between ADHD and controls.** The number of annotated genes enriched in each term (Coverage), gene names within each set, and unadjusted p-values are provided.

| ID | Ontology | Term | Coverage | p-value | Genes in set |
|---|---|---|---|---|---|
| *GO database* | | | | | |
| GO:0061046 | BP | regulation of branching involved in lung morphogenesis | 2/6 | 0.001 | *BMP4*, *SOX9* |
| GO:0072197 | BP | ureter morphogenesis | 2/7 | 0.002 | *BMP4*, *SOX9* |
| GO:1901723 | BP | negative regulation of cell proliferation involved in kidney development | 2/6 | 0.002 | *FLCN*, *BMP4* |
| GO:0032051 | MF | clathrin light chain binding | 2/7 | 0.003 | *NSG1*, *HIP1* |
| GO:0060433 | BP | bronchus development | 2/11 | 0.004 | *BMP4, SOX9* |
| GO:0060502 | BP | epithelial cell proliferation involved in lung morphogenesis | 2/10 | 0.004 | *BMP4, SOX9* |
| GO:0141176 | BP | gene silencing by piRNA-directed DNA methylation | 2/13 | 0.01 | *DNMT3A*, *MOV10L1* |
| GO:0141196 | BP | retrotransposon silencing by piRNA-directed DNA methylation | 2/13 | 0.01 | *DNMT3A*, *MOV10L1* |
| GO:0097157 | MF | pre-mRNA intronic binding | 2/13 | 0.01 | *RBM20*, *SOX9* |
| GO:0014029 | BP | neural crest formation | 2/15 | 0.01 | *SOX9*, *TCOF1* |
| GO:0140966 | BP | piRNA-mediated heterochromatin formation | 2/15 | 0.01 | *DNMT3A*, *MOV10L1* |
| GO:0098954 | CC | presynaptic endosome membrane | 1/1 | 0.01 | *VTI1B* |
| GO:0046322 | BP | negative regulation of fatty acid oxidation | 2/14 | 0.01 | *FMO4*, *SOX9* |
| GO:1903259 | BP | exon-exon junction complex disassembly | 1/1 | 0.01 | *PYM1* |
| GO:0017060 | MF | 3-galactosyl-N-acetylglucosaminide 4-alpha-L-fucosyltransferase activity | 1/2 | 0.01 | *FUT5* |
| GO:0006044 | BP | N-acetylglucosamine metabolic process | 2/16 | 0.01 | *CHST6*, *NAGK* |
| GO:0003180 | BP | aortic valve morphogenesis | 3/39 | 0.01 | *NFATC1*, *BMP4*, *SOX9* |

| | | | | | |
|---|---|---|---|---|---|
| GO:0141005 | BP | retrotransposon silencing by heterochromatin formation | 2/18 | 0.01 | *DNMT3A*, *MOV10L1* |
| GO:0018149 | BP | peptide cross-linking | 2/25 | 0.01 | *SPRR1B, TGM4* |
| GO:0048390 | BP | intermediate mesoderm morphogenesis | 1/1 | 0.01 | *BMP4* |
| *KEGG pathway database* | | | | | |
| hsa05235 | | PD-L1 expression and PD-1 checkpoint pathway in cancer | 4/87 | 0.01 | *CSNK2A2*, *JAK2*, *LCK*, *NFATC1* |
| hsa04933 | | AGE-RAGE signaling pathway in diabetic complications | 4/95 | 0.02 | *COL4A2*, *JAK2*, *NFATC1*, *PRKCZ* |
| hsa00270 | | Cysteine and methionine metabolism | 2/50 | 0.06 | *DNMT3A*, *PHGDH* |
| hsa04658 | | Th1 and Th2 cell differentiation | 3/88 | 0.06 | *JAK2*, *LCK*, *NFATC1* |
| hsa03008 | | Ribosome biogenesis in eukaryotes | 2/69 | 0.07 | *CSNK2A2*, *TCOF1* |
| hsa00920 | | Sulfur metabolism | 1/10 | 0.08 | *SQOR* |
| hsa04659 | | Th17 cell differentiation | 3/103 | 0.08 | *JAK2*, *LCK*, *NFATC1* |
| hsa00430 | | Taurine and hypotaurine metabolism | 1/16 | 0.11 | *FMO4* |
| hsa00533 | | Glycosaminoglycan biosynthesis - keratan sulfate | 1/14 | 0.13 | *CHST6* |
| hsa03013 | | Nucleocytoplasmic transport | 2/101 | 0.16 | *TMEM33*, *PYM1* |
| hsa04115 | | p53 signaling pathway | 2/75 | 0.17 | *TP53AIP1*, *COP1* |
| hsa04977 | | Vitamin digestion and absorption | 1/24 | 0.18 | *ABCC1* |
| hsa04981 | | Folate transport and metabolism | 1/30 | 0.19 | *ABCC1* |
| hsa00534 | | Glycosaminoglycan biosynthesis - heparan sulfate / heparin | 1/23 | 0.19 | *NDST1* |
| hsa00532 | | Glycosaminoglycan biosynthesis - chondroitin sulfate / dermatan sulfate | 1/20 | 0.19 | *CHPF2* |
| hsa04148 | | Efferocytosis | 3/152 | 0.20 | *DNMT3A*, *DUSP8*, *JAK2*, *NFATC1* |
| hsa00601 | | Glycosphingolipid biosynthesis - lacto and neolacto series | 1/28 | 0.20 | *FUT5* |
| hsa04064 | | NF-kappa B signaling pathway | 2/92 | 0.20 | *CSNK2A2*, *LCK* |
| hsa00260 | | Glycine, serine and threonine metabolism | 1/35 | 0.20 | *PHGDH* |

| | | | | |
|---|---|---|---|---|
| hsa00982 | Drug metabolism - cytochrome P450 | 1/57 | 0.21 | *FMO4* |

**Supplementary Table 6. Top 20 ranked GO and KEGG pathway terms linked to differentially methylated probes between diagnostic groups (remitted ADHD, persistent ADHD and controls), surviving FDR correction.** The number of annotated genes enriched in each term (Coverage), gene names within each set, and unadjusted p-values are provided.

| ID | Ontology | Term | Coverage | p-value | Genes in set |
|---|---|---|---|---|---|
| *GO database* | | | | | |
| GO:0034317 | MF | nicotinate riboside kinase activity | 1/2 | 0.001 | *NMRK2* |
| GO:0050262 | MF | ribosylnicotinamide kinase activity | 1/2 | 0.001 | *NMRK2* |
| GO:0061769 | MF | ribosylnicotinate kinase activity | 1/2 | 0.001 | *NMRK2* |
| GO:0000166 | MF | nucleotide binding | 6/2003 | 0.001 | *AGAP1*, *NMRK2*, *ACAD9*, *HSPD1*, *HSPE1*, *MOV10L1*, *PRKCZ* |
| GO:1901265 | MF | nucleoside phosphate binding | 6/2020 | 0.001 | *AGAP1*, *NMRK2*, *ACAD9*, *HSPD1*, *HSPE1*, *MOV10L1*, *PRKCZ* |
| GO:1901363 | MF | heterocyclic compound binding | 6/2147 | 0.002 | *AGAP1*, *NMRK2*, *ACAD9*, *HSPD1*, *HSPE1*, *MOV10L1*, *PRKCZ* |
| GO:0070991 | MF | medium-chain fatty acyl-CoA dehydrogenase activity | 1/3 | 0.002 | *ACAD9* |
| GO:0004466 | MF | long-chain fatty acyl-CoA dehydrogenase activity | 1/4 | 0.002 | *ACAD9* |
| GO:0043168 | MF | anion binding | 6/2276 | 0.002 | *AGAP1*, *NMRK2*, *ACAD9*, *HSPD1*, *HSPE1*, *MOV10L1*, *PRKCZ* |
| GO:0035639 | MF | purine ribonucleoside triphosphate binding | 5/1713 | 0.005 | *AGAP1*, *NMRK2*, *HSPD1*, *HSPE1*, *MOV10L1*, *PRKCZ* |
| GO:0032555 | MF | purine ribonucleotide binding | 5/1760 | 0.005 | *AGAP1*, *NMRK2*, *HSPD1*, *HSPE1*, *MOV10L1*, *PRKCZ* |
| GO:0045179 | CC | apical cortex | 1/7 | 0.005 | *PRKCZ* |
| GO:0032553 | MF | ribonucleotide binding | 5/1777 | 0.006 | *AGAP1*, *NMRK2*, *HSPD1*, *HSPE1*, *MOV10L1*, *PRKCZ* |
| GO:0071546 | CC | pi-body | 1/10 | 0.006 | *MOV10L1* |
| GO:0035748 | CC | myelin sheath abaxonal region | 1/6 | 0.006 | *PRKCZ* |
| GO:0043203 | CC | axon hillock | 1/7 | 0.006 | *PRKCZ* |

| | | | | | |
|---|---|---|---|---|---|
| GO:0120157 | CC | PAR polarity complex | 1/6 | 0.006 | *PRKCZ* |
| GO:0017076 | MF | purine nucleotide binding | 5/1853 | 0.007 | *AGAP1*, *NMRK2*, *HSPD1*, *HSPE1*, *MOV10L1*, *PRKCZ* |
| GO:0051791 | BP | medium-chain fatty acid metabolic process | 1/13 | 0.007 | *ACAD9* |
| GO:0045630 | BP | positive regulation of T-helper 2 cell differentiation | 1/7 | 0.007 | *PRKCZ* |
| *KEGG pathway database* | | | | | |
| hsa04062 | | Chemokine signaling pathway | 2/185 | 0.01 | *PRKCZ*, *CCL17* |
| hsa04144 | | Endocytosis | 2/241 | 0.01 | *AGAP1*, *PRKCZ* |
| hsa00760 | | Nicotinate and nicotinamide metabolism | 1/37 | 0.02 | *NMRK2* |
| hsa04061 | | Viral protein interaction with cytokine and cytokine receptor | 1/93 | 0.03 | *CCL17* |
| hsa04930 | | Type II diabetes mellitus | 1/45 | 0.04 | *PRKCZ* |
| hsa04657 | | IL-17 signaling pathway | 1/89 | 0.04 | *CCL17* |
| hsa04625 | | C-type lectin receptor signaling pathway | 1/102 | 0.07 | *CCL17* |
| hsa04933 | | AGE-RAGE signaling pathway in diabetic complications | 1/95 | 0.07 | *PRKCZ* |
| hsa04931 | | Insulin resistance | 1/105 | 0.08 | *PRKCZ* |
| hsa05418 | | Fluid shear stress and atherosclerosis | 1/132 | 0.08 | *PRKCZ* |
| hsa04071 | | Sphingolipid signaling pathway | 1/120 | 0.09 | *PRKCZ* |
| hsa04926 | | Relaxin signaling pathway | 1/125 | 0.09 | *PRKCZ* |
| hsa04910 | | Insulin signaling pathway | 1/131 | 0.10 | *PRKCZ* |
| hsa04611 | | Platelet activation | 1/124 | 0.10 | *PRKCZ* |
| hsa04060 | | Cytokine-cytokine receptor interaction | 1/263 | 0.11 | *CCL17* |
| hsa05415 | | Diabetic cardiomyopathy | 1/179 | 0.11 | *PRKCZ* |
| hsa04530 | | Tight junction | 1/162 | 0.11 | *PRKCZ* |
| hsa04390 | | Hippo signaling pathway | 1/154 | 0.12 | *PRKCZ* |
| hsa04360 | | Axon guidance | 1/175 | 0.15 | *PRKCZ* |
| hsa04015 | | Rap1 signaling pathway | 1/206 | 0.16 | *PRKCZ* |
